# Cryo-EM structures of human TRPC5 reveal interaction of a xanthine-based TRPC1/4/5 inhibitor with a conserved lipid binding site

**DOI:** 10.1101/2020.04.17.047456

**Authors:** David J. Wright, Katie J. Simmons, Rachel M. Johnson, David J. Beech, Stephen P. Muench, Robin S. Bon

## Abstract

TRPC1/4/5 channels are non-specific cation channels implicated in a wide variety of diseases, and TRPC1/4/5 inhibitors have recently entered the first clinical trials. However, fundamental and translational studies require a better understanding of TRPC1/4/5 channel regulation by endogenous and exogenous factors. Although several potent and selective TRPC1/4/5 modulators have been reported, the paucity of mechanistic insights into their modes-of-action remains a barrier to the development of new chemical probes and drug candidates. The xanthine class of modulators includes the most potent and selective TRPC1/4/5 inhibitors described to date, as well as TRPC5 activators. Our previous studies suggest that xanthines interact with a, so far, elusive pocket of TRPC1/4/5 channels that is essential to channel gating. Targeting this pocket may be a promising strategy for TRPC1/4/5 drug discovery. Here we report the first structure of a small molecule-bound TRPC1/4/5 channel – human TRPC5 in complex with the xanthine Pico145 – to 3.0 Å. We found that Pico145 binds to a conserved lipid binding site of TRPC5, where it displaces a bound phospholipid. Our findings explain the mode-of-action of xanthine-based TRPC1/4/5 modulators, and suggest a structural basis for TRPC1/4/5 modulation by endogenous factors such as (phospho)lipids and Zn^2+^ ions. These studies lay the foundations for the structure-based design of new generations of TRPC1/4/5 modulators.

## Introduction

Transient Receptor Potential Canonical (TRPC) proteins form homo- or heterotetrameric, non-selective cation channels permeable by Na^+^ and Ca^2+^.^1–6^ Based on sequence similarity, the mammalian TRPC proteins can be further divided into sub-groups: <u>TRPC1</u>/<u>4</u>/<u>5</u>, <u>TRPC3</u>/<u>6</u>/<u>7</u> and <u>TRPC2</u>,^7^ the latter of which is encoded by a pseudogene in humans.^8^ TRPC4 and TRPC5 are the most closely related TRPC proteins^9^ (70% sequence identity) and can form functional, homomeric TRPC4:C4 and TRPC5:C5 channels.^10^ TRPC1 (48% and 47% sequence identity to TRPC4 and TRPC5, respectively) is not thought to form functional homomeric channels, but is widely expressed and an important contributor to heteromeric TRPC1/4/5 ion channels.^10–15^

Although disruption of the *Trpc4/5* genes^16^ and global expression of a dominant-negative mutant TRPC5^12^ do not cause catastrophic phenotypes in rodents, TRPC1/4/5 channels have been implicated in a wide range of physiological and pathological mechanisms.^4,5,10,17^ These findings have driven the development of potent and selective TRPC1/4/5 modulators as chemical probes and drug candidates,^5,10,18^ and clinical trials have been started by Hydra Biosciences/Boehringer Ingelheim (the TRPC4/5 channel inhibitor BI 135889 for treatment of anxiety/CNS disorders) and Goldfinch Bio (the TRPC5 channel inhibitor GFB-887 for genetically-driven kidney disease).

Physiological activation and modulation of TRPC1/4/5 channel activity is complex,^4–6^ and may include mediation by endogenous and dietary lipids.^12,19–22^ In addition, structurally diverse pharmacological modulators have been reported,^5,10,18^ including inhibitors suitable for studies of TRPC1/4/5 in cells, tissues and animal models such as the xanthines <u>Picol45</u> (also called HC-608)^23–25^ and <u>HC-070</u>,^24,25^ the pyridazinone derivative GFB-8438,^26^ and the benzimidazole <u>ML204</u>.^27^ However, structural insight into the mode-of-action of small-molecule TRPC1/4/5 modulators is lacking, and no small-molecule binding sites have been identified.

Currently, the most potent and selective TRPC1/4/5 inhibitor is the xanthine Pico145, which inhibits the channels with IC50 values in the picomolar to low nanomolar range and displays the highest potency against heteromeric channels.^23^ Pico145 is orally bioavailable and has been used successfully for studies of TRPC1/4/5 channels in vivo.^25,28,29^ Detailed characterisation of Pico145 and the related xanthine AM237 - a partial TRPC5 agonist that inhibits other TRPC1/4/5 channels^30^ – revealed that: 1) Pico145 and AM237 act rapidly and reversibly in outside-out excised patch recordings, highlighting a membrane-delimited effect;^23,30^ 2) Pico145 is a competitive antagonist of <u>(-)-englerin A</u> (EA)^23^ and <u>AM237</u>,^30^ and also inhibits channel activation by <u>sphingosine-1-phosphate</u> (S1P), <u>carbachol</u> and <u>Gd^3+^</u>,^23,24^ although picomolar concentrations of Pico145 can potentiate Gd^3+^-induced TRPC4 currents as well;^23^ 3) Pico145 concentration-dependently inhibits photoaffinity labelling of TRPC5 by the xanthine-based photoaffinity probes Pico145-DAAlk and Pico145-DAAlk2, with IC50 values in the same range as inhibition of TRPC5-mediated calcium influx.^31^ These results provide evidence that these xanthines modulate TRPC1/4/5 channels through a direct molecular interaction with the channels and are consistent with the hypothesis that Pico145 and AM237 bind to a well-defined, high-affinity binding site essential to TRPC1/4/5 channel gating. Targeting this binding site may be a promising strategy to develop further generations of potent and selective TRPC1/4/5 chemical probes and drug candidates.

Several high resolution TRPC structures have been determined by single-particle cryo-electron microscopy (cryo-EM), including zebrafish TRPC4,^32^ mouse TRPC4,^33^ and mouse TRPC5.^34^ To date, <u>TRPC6</u> is the only TRPC channel for which small molecule-bound structures have been reported.^35,36^ Here, we report the first structure of a TRPC1/4/5 channel in complex with a small-molecule modulator: the homomeric TRPC5:C5 channel in complex with Pico145. We found that Pico145 binds between the trans-membrane domains of two TRPC5 subunits, where it displaces a phospholipid in a conserved lipid binding site. Docking and mutagenesis studies support these findings, and provide additional insights into the mode-of-action of xanthines as modulators of TRPC5 and TRPC4 channels. In addition, we have identified a putative intracellular zinc binding site of TRPC5 that is conserved within the TRPC family. This work provides a rational basis for the design of new TRPC1/4/5 modulators and suggests important functional roles for the conserved TRPC1/4/5 lipid and zinc binding sites. Therefore, our findings may help unravel physiological mechanisms of TRPC1/4/5 modulation, and pave the way for structure-based TRPC1/4/5 drug discovery efforts.

## Results

### Functional characterisation of C-terminally truncated human TRPC5

For our structural studies, we engineered a human TRPC5 construct containing an N-terminal maltose binding protein (MBP), followed by a PreScission protease site (PreS) and a C-terminally truncated (Δ766-975) hTRPC5 (99% identical to the analogous mTRPC5 construct^34^). Upon over-expression in HEK293 cells, TRPC5:C5 channels formed by this construct were activated by EA (EC50 3 nM; cf. EC50 2 nM for full-length hTRPC5) and inhibited by Pico145 (IC50 4 nM; cf. IC50 2 nM for full-length hTRPC5) **(Supplementary Figure 1)**. These results suggest MBP-PreS-hTRPC5_Δ766-975_ as a suitable construct for structural determination of the xanthine binding site.

### Protein purification and quality control

Baculoviruses were made for MBP-PreS-hTRPC5_Δ766-975_ expression and used to transfect Freestyle^™^ 239-F Cells in suspension. MBP-tagged TRPC5 was purified using a protocol adapted from Duan et al.,^34^ but with amphipol (PMAL-C8) exchange performed on-resin and ultracentrifugation used in lieu of size exclusion chromatography to remove aggregated material. The protein was pure according to SDS-PAGE analysis, and negative stain electron microscopy showed a monodisperse sample with particles consistent in size with the expected tetrameric structure (**Supplementary Figure 2**). Further negative stain analysis of the TRPC5 protein in the presence of Pico145 showed a similar monodispersity with no evidence for any gross conformational changes or disruption of the tetramer.

### Structure of a human TRPC5:C5 channel in complex with Pico145

The cryo-EM structure of a human TRPC5:C5 channel in the presence of Pico145 was determined with a global resolution of 3.0 Å using C4 symmetry (**Figure 1**; **Supplementary Table 1**; **Supplementary Figure 3**). To investigate if there was heterogeneity between the 4 subunits, the complex was also refined with no symmetry imposed (C1 symmetry). The resulting TRPC5 structure showed no significant difference to the C4 refined model, but the global resolution was lower (3.3 Å). The tetrameric structure of TRPC5 was built (based on the mTRPC5 *apo* structure, PDB 6aei) and showed the typical TRPC fold: a large intracellular domain made up of the N- and C-termini of each monomer, six transmembrane helices per monomer and relatively short extracellular loops (**Figure 1B**). The N-terminus (residues 1-366) folds into 4 ankyrin repeats, a helical linker domain and the pre S1-elbow domain. The first four transmembrane helices fold into a voltage sensing-like domain (VSLD); the next two helices form the pore domain, together with the re-entrant pore helix (E3 loop), which lines the ion permeation pathway. The C-terminus (residues 626 onwards) contains the TRP domain and the coiled-coil domain, which is formed by one helix from each monomer. The maltose binding protein (MBP), protease site (PreS) and the first 16 amino acid residues of TRPC5 were not resolved in the map, likely because of flexibility. Likewise, residues 37-60, 74-95, 118-134, 273-285, 387-391, 665-702 and 759-765 were not modelled due to them being poorly resolved. We tentatively modelled residues 734 to 759, yet the position of these residues should be interpreted with caution. All residues that were not modelled are found in the intracellular domain or extracellular loop 1; the transmembrane helices and the other extracellular loops were modelled into the electron density.

**Figure 1:**
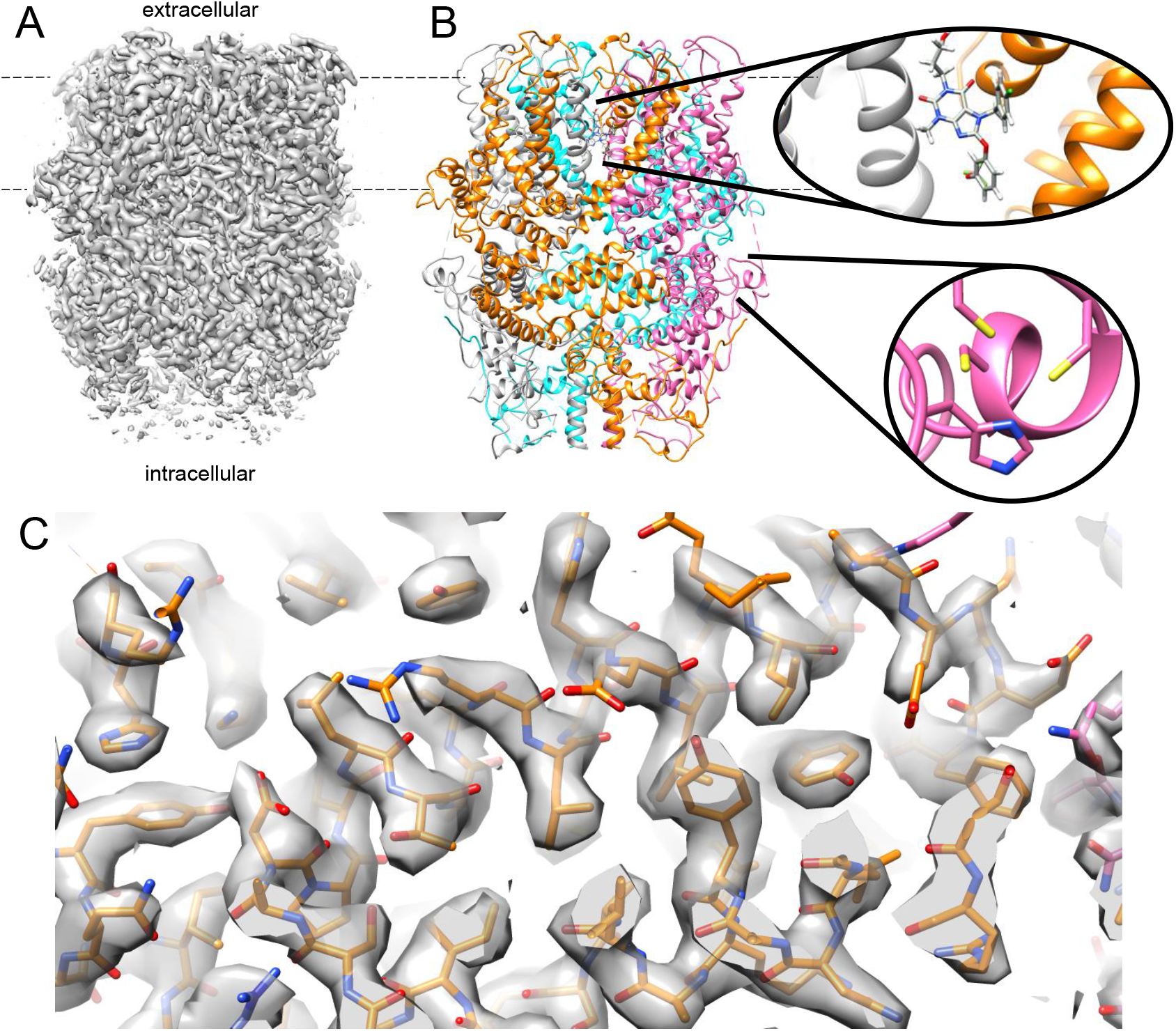
Structure of human TRPC5 in complex with Pico145. A) Cryo-EM density map of TRPC5:Pico145. B) Model of TRPC5:Pico145 coloured by monomer, showing the typical TRP channel domain swap architecture. Dashed lines indicate the position of the lipid bilayer. Insets show one of the four Pico145 binding sites (between trans-membrane domains of the blue and orange TRPC5 monomers) and one of the four putative intracellular zinc binding sites (in the magenta monomer). C) Representative fit of TRPC5:Pico145 model (B) into the experimental density (A) showing helical regions spanning residues 193 to 234.

Comparison of our TRPC5 structure to the previously reported 2.9 Å mTRPC5 *apo* structure^34^ revealed a backbone (Cα) RMSD of 0.82 Å showing a similar overall structure, consistent with both being in the closed state. There were, however, several notable local differences between the *apo* and Pico145-bound structures (**Supplementary Figure 4A,B**). In our structure, the coiled-coil domain was less well ordered, and residues 734 to 759 could not be fit with confidence. However, the quality of our map was such that additional residues could be built within the region 175-187, and that residues 172-174 could be placed with greater accuracy. This region showed that histidine 172 and cysteines 176, 178 and 181 all point towards a central density, consistent with metal ion binding (**Figure 1B** and **Figure 5**; see below for details). In addition, there was a notable difference in the non-protein density between the *apo* and Pico145-bound structures, as discussed below.

**Figure 2:**
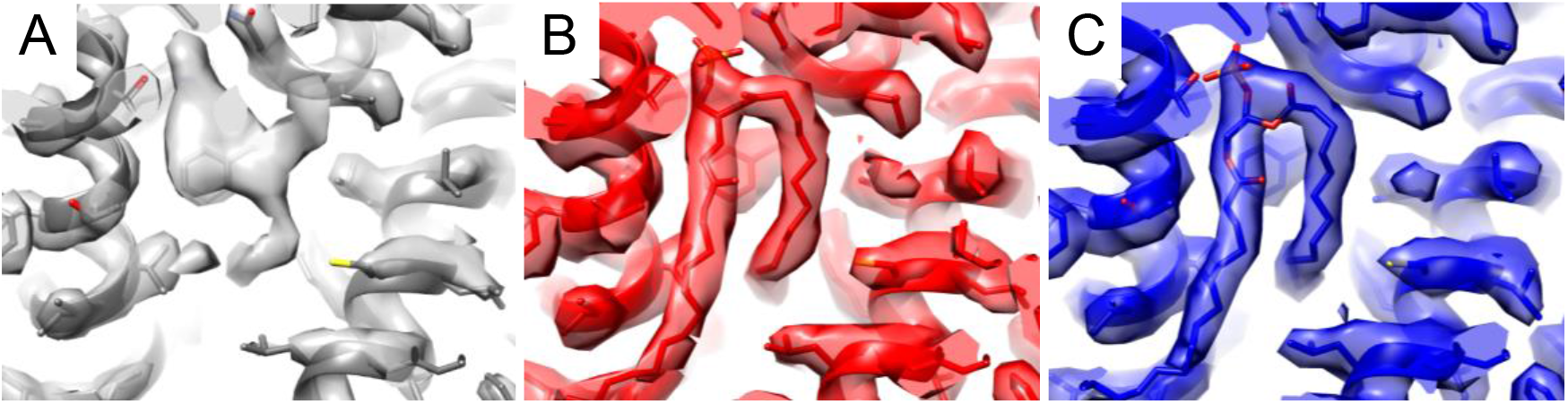
Pico145 binds to a conserved TRPC4/5 lipid binding site. A) Density observed in our hTRPC5:Pico145 structure (100 μM Pico145). B) Equivalent density in the published mTRPC5 *apo* structure (PDB 6aei, EMDB 9615) that was modelled as a phospholipid. C) The same site in the published mTRPC4 structure (PDB 5z96, EMDB 6901).

**Figure 3:**
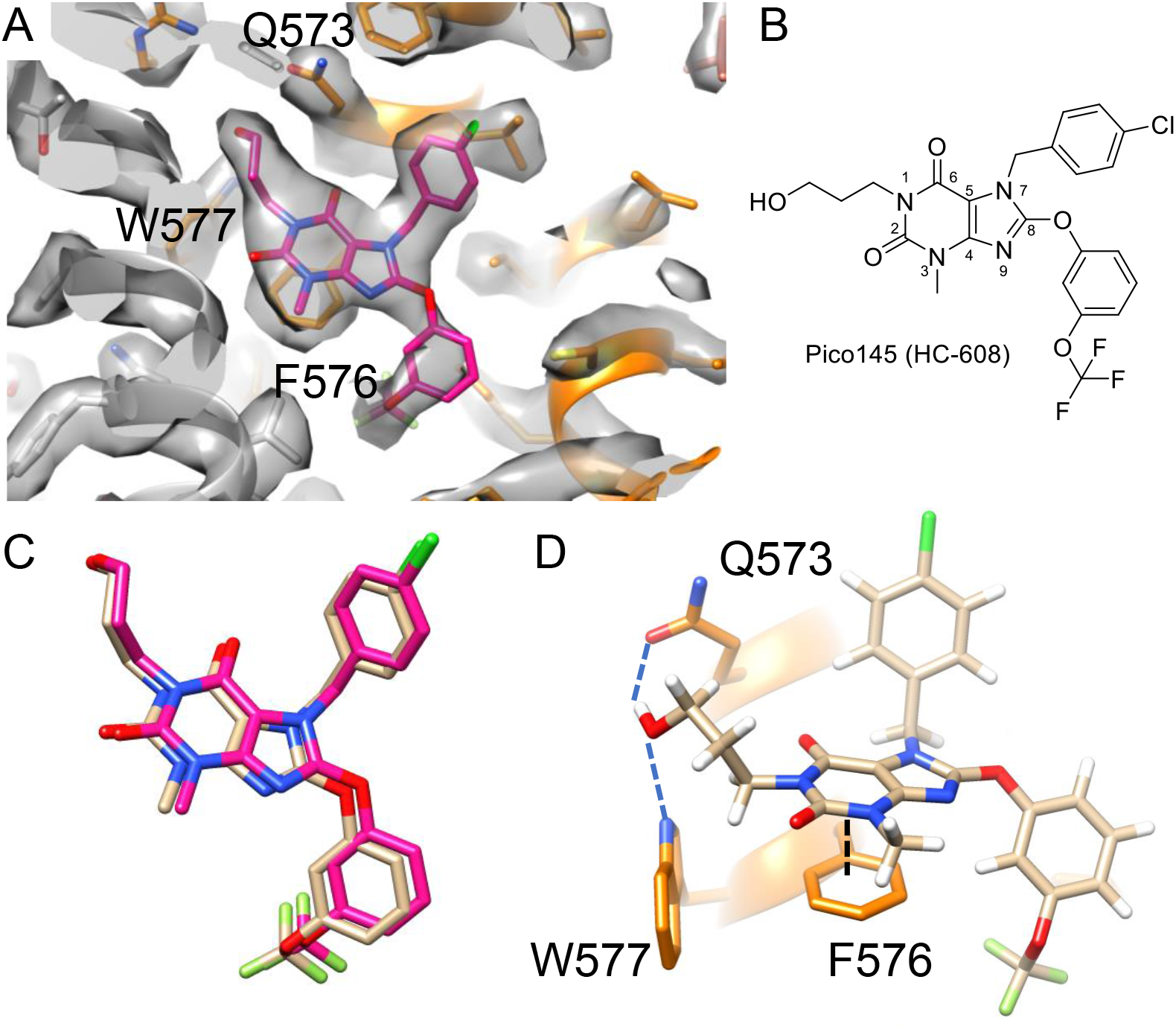
Analysis of the Pico145 binding site of TRPC5. A) The refined structure of the Pico145 binding site, showing the electron density map determined by cryo-EM (grey) and fitted Pico145 (magenta). The binding site spans two monomers: monomer 1 is shown in orange and monomer 2 is shown in grey. Three of the key residues for interaction, Q573, F576 and W577, are indicated. B) Chemical structure of Pico145 with numbering of the xanthine core. C) An overlay of the top docking pose of Pico145 (peach) and our fitted Pico145 molecule (magenta). The top 3 docking poses are shown in **Supplementary Figure 7**. D) The top scoring docking pose suggests that Pico145 hydrogen bonds to Q573 and W577 (dashed blue lines) and π-stacks with F576 (dashed black line).

**Figure 4:**
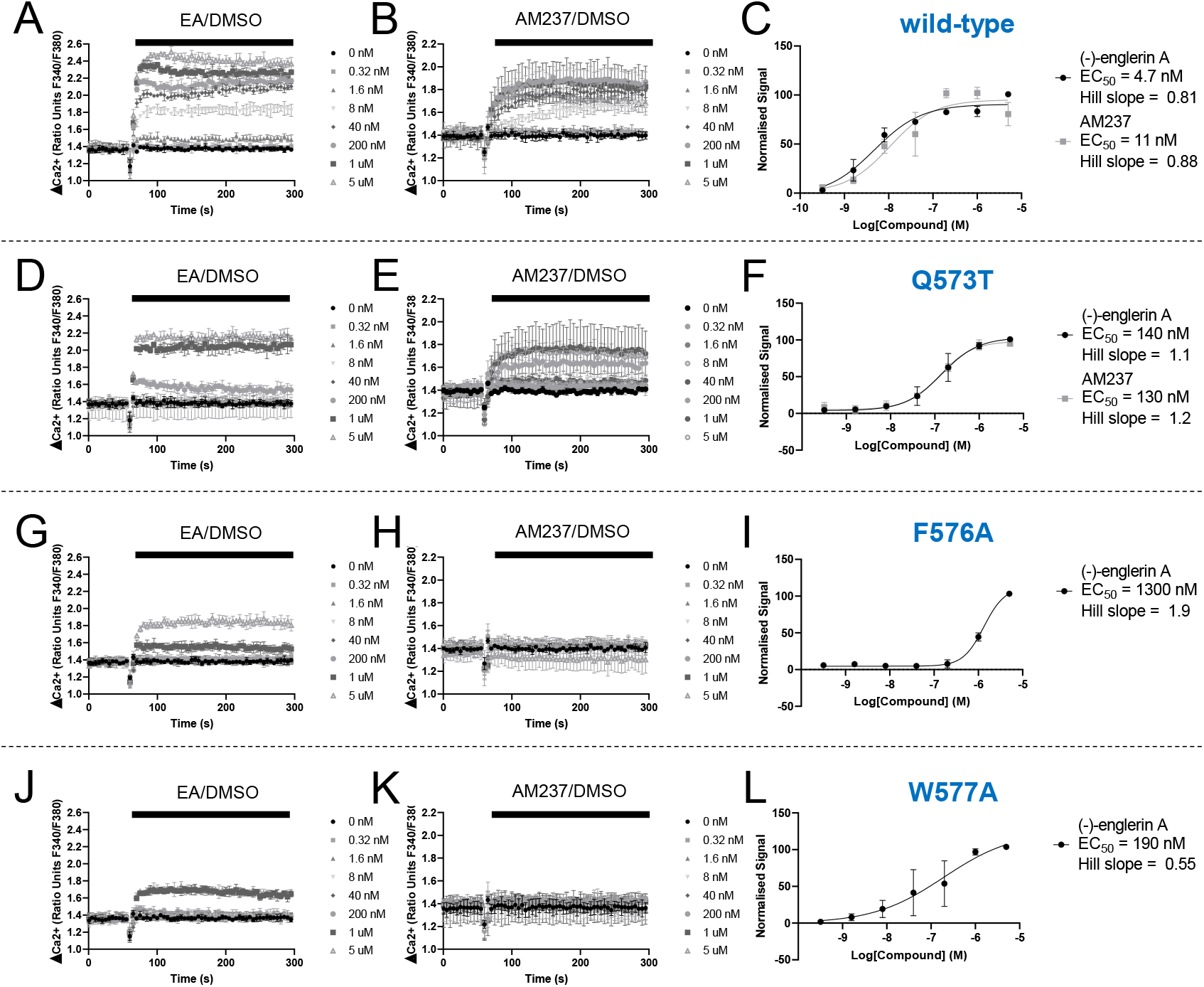
Mutations in the xanthine binding site of TRPC5 affect responses to the xanthine-based TRPC5 agonist AM237. A,B,D,E,G,H,J,K) Representative traces from single 96-well plates (N = 3) showing an increase in [Ca^2+^]_i_ in response to 0.32 nM to 5 μM of EA (A, D, G, J) or AM237 (B, E, H, K) in HEK 293 cells transiently expressing wild-type TRPC5-SYFP2 (A,B), TRPC5-SYFP2_Q573T_ (D,E), TRPC5-SYFP2_F576A_ (G,H) or TRPC5-SYFP2_W577A_ (J,K). C) Concentration-response data for experiments in (A) and (B) (mean normalised response ± SEM; n/N = 3/6). F) Concentration-response data for experiments in (D) (mean normalised response ± SEM; n/N = 3/6) and (E) (mean normalised response ± SEM; n/N = 2/4). I) Concentration-response data for experiments in (G) (mean normalised response ± SEM; n/N = 2/4; AM237 gave no response up to 5 μM). L) Concentration-response data for experiments in (J) (mean normalised response ± SEM; n/N = 2/5; AM237 gave no response up to 5 μM). Responses were calculated at 240-300 s compared to [Ca^2+^]_i_ at baseline (0-60 s). Averages of technical repeats of each individual experiment were normalised to the maximum response for each set of experiments, combined, and fit with GraphPad Prism 8 (variable slope, four-parameter fit).

**Figure 5:**
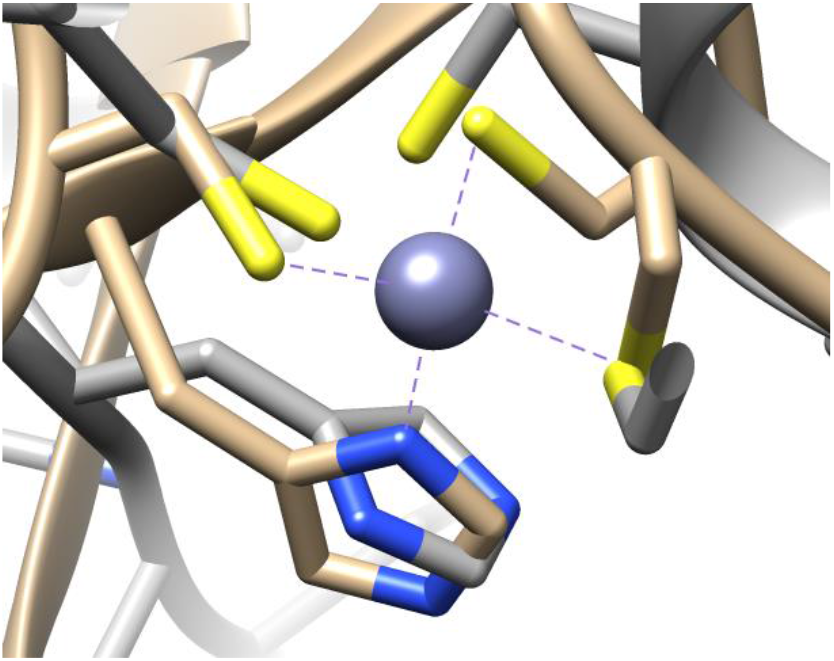
Structure alignment of hTRPC5 and hURF1 is consistent with the identification of a TRPC5 zinc binding site. Sequence alignment revealed that the Cys3-His1 motif closely matched that of the PHD finger of human UHRF1 (an unrelated zinc-binding protein). Manual alignment was performed between hTRPC5 (grey) and hUHRF1 (PDB 3zvz; wheat).

### Pico145 binds to a conserved lipid binding site of TRPC5

Comparison of the cryo-EM structure of TRPC5:Pico145 to published mTRPC5^34^ and mTRPC4^33^ *apo* structures showed clear differences in one of the lipid binding sites (**Figure 2A-C**). In the mTRPC5 *apo* structure, a density with a characteristic lipid U-shape was observed close to the E3 re-entrant loop (**Figure 2B**). This density was ascribed to a phospholipid (postulated to be a phosphatidic acid or ceramide-1-phosphate)^34^ that interacts with the LFW motif, which is conserved in the TRPC family (**Supplementary Table 2**). Indeed, a similar density is present in the mTRPC4 structure (**Figure 2C**). The hTRPC3^35,37^ and hTRPC6^35,36^ structures contain lipids that interact with their LFW motifs as well, but with altered geometry (**Supplementary Figure 5**). Our TRPC5:Pico145 structure also showed density in this region. However, the shape of this density was strikingly different to that in the mTRPC5 and mTRPC4 *apo* structures (**Figure 2A-C**): in our structure, no density was observed for one of the lipid tails, and additional density was found between residues Q573, F569 and L572. Although this density was not consistent with a bound lipid, its shape and size were consistent with the presence of a molecule of Pico145. Indeed, modelling of Pico145 into this density gave an excellent fit (**Figure 3A**). Similar to the phospholipid in the mTRPC5 and mTRPC4 *apo* structure, Pico145 (**Figure 3B**) was observed between two TRPC5 monomers, with four molecules of Pico145 per TRPC5 tetramer. This was observed in both the C4 and C1 reconstructions and is consistent with all four sites being occupied.

Fitting Pico145 into the density shows that most of the TRPC5 residues interacting with the inhibitor are from helix S5 and the pore helix of the monomer on the right (monomer 1; **Figure 3a**; orange). The 3-hydroxypropyl substituent on N-1 of Pico145 interacts with the side chains of W577 and Q573, apparently substituting for the phospholipid phosphate group in the *apo* structures. Pico145’s xanthine core π-stacks with F576, and several residues (including L521, Y524, C525 and V579) interact with the 3-(trifluoromethoxy)phenoxy substituent on C-8 of Pico145. The 4-chlorobenzyl substituent on N-7 of Pico145 points upwards, away from the lipid site, making a non-polar interaction with L572. Pico145 also makes additional interactions with helix S6 of the monomer on the left (monomer 2; **Figure 3a**; grey): Pico145 interacts with T603 and packs against G606, and several hydrophobic amino acids (V610 and V614) interact with Pico145’s 3-(trifluoromethoxy)phenoxy substituent. It should be noted that the local resolution of the cryo-EM map at the core of the TRPC5 channel, including the Pico145 binding site, was better than 2.9 Å, making the fit unambiguous **(Supplementary Figure 3**).

During the optimisation of conditions for TRPC5:Pico145 cryo-grid preparation, we varied concentrations of TRPC5 and Pico145. As part of this process, an additional structure of TRPC5 in the presence of Pico145 was solved with TRPC5 at 1 mg·ml^-1^ and Pico145 at 50 μM (compared to 2 mg·ml^-1^ TRPC5 and 100 μM Pico145 in the structure described above) to a global resolution of 2.9 Å. The density around the Pico145 binding site in this structure showed features consistent with both the bound phospholipid and Pico145, suggesting partial occupancy under these conditions (**Supplementary Figure 6A-C**). These data further support the observation that Pico145 can displace each of the four phospholipids bound to a tetrameric TRPC5 channel.

### Docking of xanthines into TRPC5 and TRPC4 channel structures

To further investigate the Pico145 binding site of TRPC5 we conducted docking experiments, using our TRPC5:Pico145 cryo-EM structure in which the coordinates for the Pico145 were removed. Docking of Pico145 into a 36×36×36 Å grid around the Pico145 binding site produced three poses with similar predicted binding energies. These poses are almost identical to each other (**Supplementary Figure 7A**), and to the pose of Pico145 modelled in our cryo-EM structure (**Figure 3C**), giving confidence in the docking procedure and further validation to the observed density being assigned to Pico145. The lowest energy pose suggests that the 3-hydroxypropyl substituent at N-1 of Pico145 makes two hydrogen bonds, donating a hydrogen bond to the carbonyl of the side chain of Q573 and accepting a hydrogen bond from the indole of W577 (**Figure 3D**, **Supplementary Figure 7B**). This 3-hydroxypropyl substituent occupies a position similar to the phosphate head group of the phospholipid in other structures. Consistent with our TRPC5:Pico145 structure, all three poses show a π-stacking interaction between F576 and the xanthine core of Pico145 (**Figure 3D, Supplementary Figure 7**). Although the positively charged R557 is nearby, no significant interactions with this residue appear to take place. The xanthine core of Pico145 and the hydrophobic tails overlay almost perfectly in all three docked poses (**Supplementary Figure 7A**). The main difference between poses is in the conformation of the 3-hydroxypropyl substituent, suggesting it is flexible in the TRPC5 binding pocket, where it may form hydrogen bonds with different TRPC5 residues. Structural alteration of the docked Pico145 in TRPC5 to the related xanthine AM237 (by addition of a chlorine atom) resulted in a clash with L521 (**Supplementary Figure 8**), suggesting a structural rearrangement is required for AM237 to bind to TRPC5. This could explain why Pico145 is an inhibitor of TRPC5, whereas AM237 is a partial agonist.^30^

To gain insight into the potent inhibition of TRPC4 channels by Pico145,^23^ we next docked Pico145 into the published mTRPC4 *apo* structure (PDB 5z96; the phospholipid was removed before docking). The docking suggests that Pico145 adopts a similar pose in TRPC4 and TRPC5, making similar interactions with conserved residues, particularly residues equivalent to the TRPC5 residues Q573, F576 and W577 (**Supplementary Figure 9**). These data suggest that the binding site of Pico145 is conserved between TRPC4 and TRPC5 channels.

### Mutations in the xanthine binding site alter the response of TRPC5 to AM237

TRPC4 and TRPC5 channels (including heteromeric TRPC1/4/5 channels containing TRPC1) are modulated by the xanthines Pico145 and AM237, whereas TRPC3/6/7 channels are not.^23,24^ This is despite the high sequence similarity between TRPC proteins around the xanthine/lipid binding sites of TRPC4 and TRPC5 channels (**Supplementary Table 2**). According to our structural modelling and docking, several residues were predicted to be important for binding of xanthines to TRPC5, and we decided to make and test three TRPC5 variants: Q573T, F576A and W577A (**Figure 3D, Table 1**). Q573 is conserved between TRPC4 and TRPC5, but the analogous residues in TRPC1 and TRPC3/6/7 are phenylalanine and lysine, respectively (**Supplementary Table 2**). Q573T was chosen because threonine and glutamine are both hydrophilic amino acids with the propensity to form hydrogen bonds, but with altered geometries, which might affect xanthine binding. F576 and W577 are fully conserved TRPC residues that appear to interact with the co-purified lipid in TRPC3,^35,37^ TRPC6,^35,36^ TRPC4^33^ and TRPC5^34^ structures (**Supplementary Figure 5**), and therefore could be important for function or molecular interactions. F576A should still be sufficiently hydrophobic to be stable in the membrane environment, but would not be able to π-stack with the xanthine core of Pico145/AM237. W577A would also retain the hydrophobic character at this position, but would lose the ability of the W577 indole NH to act as a hydrogen bond donor to the 3-hydroxypropyl group of Pico145/AM237.

**Table 3:** Summary of data in **Figure 4**.

| TRPC5 variant | Residue in TRPC4/1/3/6/7 | EA EC<sub>50</sub> (nM) | AM237 EC<sub>50</sub> (nM) |
| --- | --- | --- | --- |
| Wild-type |  | 4.7 | 11 |
| Q573T | Q/F/K/K/K | 140 | 130 |
| F576A | F/F/F/F/F | 1300 | No response up to 5 μM |
| W577A | W/W/W/W/W | 190 | No response up to 5 μM |

The constructs were tested using fluorometric intracellular calcium ([Ca^2+^]_i_) measurements (**Figure 4**). Activation by EA^11^ was used to show that TRPC5 variants were expressed and formed functional TRPC5:C5 channels. The inhibitory potency of Pico145 on TRPC1/4/5 channels is dependent on the EA concentration used in assays,^23^ which was expected to complicate analysis of Pico145 responses of TRPC5 variants. Therefore, we instead tested TRPC5 responses to AM237, which is thought to occupy the same binding site as Pico145 (see above and ^30,31^). Mutations that resulted in TRPC5 activation by EA, but not by AM237, were likely to reveal residues contributing to xanthine binding. Wild-type TRPC5 was activated by EA (EC_50_ 4.7 nM) and AM237 (EC_50_ 11 nM) (**Figure 4A-C**). Both EA (EC_50_ 140 nM) and AM237 (EC_50_ 130 nM) activated the TRPC5_Q573T_ variant (**Figure 4D-F**), but potencies of both activators were lower by more than an order of magnitude compared to wild-type TRPC5. TRPC5_F576A_ (**Figure 4G-I**) and TRPC5_W577A_ (**Figure 4J-L**) could be activated by EA, but again much higher EA concentrations were needed (EC_50_ values of 1300 and 190 nM, respectively). However, neither TRPC5 variant could be activated by AM237 (no response up to 5 μM AM237), suggesting that that F576 and W577 are key residues for the binding/gating of TRPC5 by AM237. These results are consistent with our structural data and modelling/docking.

### The TRPC5:Pico145 structure reveals a putative, conserved zinc binding site

Our TRPC5 cryo-EM map allowed residues 172-187 to be resolved and built. This region, which was not built into the previously published mTRPC5 *apo* structure,^34^ includes residues H172, C176, C178 and C181. The nature of the amino acids and the directionality of their thiol and imidazole substituents suggested the presence of a metal binding site. A PDB motif query search, using hTRPC5 residues 172-181 (HXXXCXCXXC), revealed the presence of a similar binding site in the PHD domain of the zinc-binding E3 ubiquitin ligase UHRF1 (PDB 3zvz) where the motif coordinates a Zn^2+^ ion. Manual overlay of this structure with our TRPC5 structure revealed a close alignment of residues (**Figure 5**). The importance of this site is reflected by the complete conservation of these residues in the TRPC family (**Supplementary Figure 10**). In addition, Park et al. recently reported that intracellular application of Zn^2+^ (micromolar concentrations) can open neuronal TRPC5 channels through an unknown mechanism.^38^ Therefore, we solved an additional structure of TRPC5 (1 mg·ml^-1^; ca. 7 μM of TRPC5 monomer) in the presence of ZnCl_2_ (final concentration 20 μM) (**Supplementary Figure 3C**). This structure, which was determined to a global resolution of 2.8 Å, showed that addition of ZnCl_2_ did not cause significant structural changes compared to our TRPC5:Pico145 structure described above. Again, the C-terminal coiled-coiled tail was more disordered than in the published mTRPC5 *apo* structure, suggesting that our slight alterations of the construct or purification protocol may be the cause of this disorder (**Supplementary Figure 4**). The lipid/xanthine binding site showed a non-protein density consistent with a phospholipid (**Supplementary Figure 6A,C,D**). In the zinc binding domain, the densities were similar (and partially disordered) in the TRPC5:Zn^2+^ and TRPC5:Pico145 structures, suggesting that the site may already be fully occupied in the absence of added ZnCl_2_. A structural alignment showed that the putative zinc binding motif is present in the electron density maps of all TRPC channel structures; it was built into the hTRPC3^35,37^ and hTRPC6^35,36^ structures, but not into the mTRPC4^33^ or mTRPC5^34^ structures (**Supplementary Figure 10**). These findings suggest that TRPC channels have a conserved intracellular zinc binding site.

## Discussion

The widely expressed TRPC1/4/5 channels have been implicated in various physiological and pathological mechanisms and are an emerging class of potential drug targets.^4,5,10,17^ Our study provides the first structural insight into small-molecule modulation of a TRPC1/4/5 channel. By determination of a 3.0 Å structure of the TRPC5:C5 channel in complex with the TRPC1/4/5 inhibitor Pico145, we show that Pico145 can bind to up to four lipid binding sites between TRPC5 subunits, where it displaces phospholipids that interact with the channel’s pore helices, previously proposed to be ceramide-1-phosphate or a phosphatidic acid.^34^ Pico145 could be fit unambiguously into the non-protein density between TRPC5 subunits, which was strikingly different from the density in the corresponding sites of the published mTRPC5^34^ and mTRPC4^33^ *apo* structures, and of our 2.8 Å structure of TRPC5 determined in the presence of ZnCl_2_. Our findings were further corroborated by a partially-occupied TRPC5:Pico145 structure (showing density for both Pico145 and the phospholipid), molecular docking, and site-directed mutagenesis.

The xanthines Pico145 and HC-070 are the most potent and selective TRPC1/4/5 inhibitors to date.^23,24^ However, two closely related xanthine derivatives, AM237^30^ and the photoaffinity probe Pico145-DAAlk,^31^ are partial agonists of the homomeric TRPC5:C5 channel that inhibit other TRPC1/4/5 channels. Pico145 is a competitive antagonist of AM237^30^ and concentration-dependently inhibits both activation and photoaffinity labelling of TRPC5:C5 by Pico145-DAAlk.^31^ These data are consistent with the hypothesis that xanthine-based TRPC1/4/5 modulators all bind to the same lipid binding site of TRPC1/4/5 channels, where they can stabilise either open or closed channel conformations depending on xanthine substituent patterns and the composition of the specific TRPC tetramer. Therefore, targeting the xanthine binding site of TRPC1/4/5 channels may allow the development of modulators of specific TRPC channel tetramers, a major challenge in TRPC1/4/5 pharmacology.

TRPC1/4/5 channels are modulated by endogenous and dietary lipids, but the mechanisms are complex and incompletely understood.^12,19–22^ Lipids may affect TRPC1/4/5 channels indirectly, e.g., the phospholipid S1P can activate channels through G-protein signalling.^23^ However, more direct mechanisms are possible as well, e.g., lysophosphatidylcholine (LPC) activates TRPC5 in excised outside-out patches in the absence of GTP.^39^ Such mechanisms may involve direct molecular interactions and/or changes to the membrane environment of TRPC1/4/5 channels. The TRPC5 lipid binding site that Pico145 interacts with is highly conserved within the TRPC family (**Supplementary Table 2**). Bound (phospho)lipids have been found in this site in structures of hTRPC3,^35,37^ mTRPC4,^33^ mTRPC5,^34^ and hTRPC6,^35,36^ and the TRPC6 activator AM-0883 displaces the ordered lipid bound in this site near the P-loop of hTRPC6 (**Supplementary Figure 5F**).^36^ The hTRPC5 residues F576 and W577 are conserved throughout the TRPC family and are likely to be responsible for lipid binding. Mutations of these residues resulted in TRPC5 variants that displayed lower sensitivity to EA and lacked response to AM237. This suggests that EA may bind in the lipid/xanthine binding site as well, which is consistent with the observation that the potency of Pico145 is dependent on the concentration of EA used in assays.^23^ Our study suggests that (phospho)lipids binding to the conserved TRPC lipid binding sites may have a direct functional effect on channel gating, and that this site can be targeted by TRPC1/4/5 activators as well as inhibitors. However, further studies are required to fully understand the mechanisms of TRPC1/4/5 channel binding and gating by (phospho)lipids and different small-molecule TRPC1/4/5 activators such as EA.

Intracellular Zn^2+^-dependent activation of TRPC5 channels was recently reported to contribute to oxidative neuronal death,^38^ but the molecular mechanism of Zn^2+^ regulation of TRPC5 was not identified. We have identified a putative intracellular zinc binding site of TRPC5, which is conserved in all TRPC channels. Cryo-EM of TRPC5 in the presence of ZnCl_2_ resulted in a further TRPC structure determined to a resolution of 2.8 Å, the highest resolution TRPC5 structure to date. Presence of ZnCl_2_ did not change the overall structure compared to TRPC5:Pico145. This observation may indicate that zinc is already bound in TRPC5:Pico145, or that the role of zinc in TRPC5 modulation is subtle, which is consistent with the delayed Zn^2+^-mediated [Ca^2+^]_i_ increase observed in calcium imaging experiments, and the small TRPC5 currents evoked by intracellular application of ZnCl_2_.^38^ Part of the putative zinc binding site has also been implicated in TRPC5 glutathionylation: mutation of C176, C178 or C181 prevented TRPC5 opening in response to glutathionylation.^40^ A separate report proposed that palmitoylation of C181 is required for correct trafficking of TRPC5 to the plasma membrane in striatal neurons.^41^ These data suggest that this small domain may be an important regulatory node of TRPC channels, and that further experiments are needed to fully understand its role in TRPC channel biology.

Overall, our data provide the first structural insight into TRPC1/4/5 channel modulation and suggest direct modulatory roles for (phospho)lipids and Zn^2+^ ions. These studies lay the foundations for the structure-based design of TRPC1/4/5 modulators, and may therefore support the development of new TRPC1/4/5 chemical probes and drug candidates for an increasing number of therapeutic areas.

## Materials and Methods

### Plasmids

All plasmids were cloned using NEBuilder HiFi DNA Assembly Master Mix (New England Biolabs) and mutations were made using the Q5 Site-directed mutagenesis kit (New England Biolabs). The BacMam vector was a kind gift from Professor Eric Gouaux (Vollum Institute).^42^ Briefly, human TRPC5 was cloned as a C-terminally truncated form (Δ766-975) with an N-terminal maltose binding protein tag followed by a PreScission protease cleavage site (as in ^34^). Mutations to the xanthine binding site were introduced into TRPC5-SYFP2 in pcDNA4.^30^ Bacmids were produced according to the Bac-to-Bac protocol (Invitrogen).

### Protein expression and purification

P2 virus was added to 2.0 million per ml of Freestyle™ 293-F Cells (Themofisher Scientific) in Gibco FreeStyle 293 Expression Medium (Invitrogen) at a final volume of 10% at 37 °C and 5% CO2. After 24 h, 5 mM sodium butyrate (Sigma Aldrich) was added and the temperature was lowered to 30 °C. After a further 48 h, cells were harvested by centrifugation and frozen. The protein purification protocol was adapted from Duan et al.^34^ Unless stated otherwise, all detergents (and amphipol PMAL-C8) were supplied by Generon. For a typical purification, a 200 ml cell pellet was thawed and re-suspended in 20 ml of 1% DDM, 0.1% CHS, 150 mM NaCl (Sigma Aldrich), 30 mM HEPES (Sigma Aldrich) pH 7.5, 1 mM DTT (Fisher Scientific Ltd) and protease inhibitor cocktail (Sigma Aldrich), and incubated by rotating at 4 °C for 1 h. After ultracentrifugation at 100,000 x g for 1 h at 4 °C, the supernatant was incubated with 400 μl bed volume of pre-washed amylose resin (New England Biolabs) for 18 h, rotating at 4 °C. The resin was washed with 20 ml of 0.1 % digitonin and 0.01% CHS, 150 mM NaCl, 30 mM HEPES pH 7.5, 1 mM DTT. The resin was re-suspended in 10 ml of 0.1% PMAL-C8, 150 mM NaCl, 30 mM HEPES pH 7.5, 1 mM DTT and rotated for 4 h at 4 °C. 100 mg of Biobeads (Bio-rad) were then added and incubated for 18 h, rotating at 4 °C, to remove detergents. The resin was washed with 20 ml of buffer containing no detergent (150 mM NaCl, 30 mM HEPES pH 7.5, 1 mM DTT) before eluting in 5 ml of the same buffer plus 40 mM maltose (Sigma Aldrich). The eluate was subjected to ultracentrifugation at 100,000 x g for 1 h at 4 °C to remove any precipitated material and the supernatant was concentrated to the required concentration with 100 kDa cut off vivaspin 500 concentrators (Sigma Aldrich). In a typical purification protocol, 50-100 μg of purified TRPC5 was produced from 200 ml of cell suspension.

### Negative stain electron microscopy

Sample quality was assessed by negative stain electron microscopy. Briefly, carbon-coated grids were glow-discharged for 40 s in a Pelco glow discharge unit. After charging, 3 μl of purified TRPC5 at 0.04 mg·ml^-1^ was added to the grid and the sample was stained with 1% uranyl acetate, as previously described.^43^ For Pico145-containing grids, Pico145 (10 mM in DMSO) was added to 2 mg·ml^-1^ purified TRPC5 to a final concentration of 100 μM, which was subsequently diluted 50-fold in detergent free wash buffer, resulting in a final concentration of 2 μM Pico145 and 0.04 mg·ml^-1^ TRPC5. Negative stain grids were imaged using a Technai T12 microscope fitted with a LaB6 filament operating at 120 kV with a nominal magnification of 49,000 × on a 2 k × 2 K Gatan CCD camera.

### Sample preparation and data collection

Pico145 (final concentration of 100 μM, from a 10 mM DMSO stock; final DMSO concentration 1%) was added to purified TRPC5 in PMAL-C8 at 2.0 mg·ml^-1^. A 3 μl aliquot of the sample was applied to a Quantifoil Cu R1.2/1.3, 300 mesh holey carbon grid, which had been glow discharged for 30 seconds using a Pelco glow discharge unit. An FEI Vitribot was used to blot the grids for 6 seconds (blot force 6) at 100% humidity and 4 °C before plunging into liquid ethane. The grids were loaded into an FEI Titan Krios transmission electron microscope (Astbury Biostructure Laboratory, University of Leeds) operating at 300 kV, fitted with a Gatan K2 direct electron detector operating in counting mode. Automated data collection was carried out using EPU software with a defocus range of −1 to −3 μm. A total of 3,482 micrographs were collected with a pixel size of 1.07 Å. The total dose, 75 e-·Å^-2^, was acquired by use of a dose rate of 8.64 e-·pixel^-1^·s^-1^ (7.58 e-·Å^-2^·s^-1^) across 40 frames for 10 s total exposure time. The parameters for the structures in the presence of ZnCl_2_ or 50 uM Pico145 used 1 mg·ml^-1^ final protein concentration and used the same EPU software with a defocus range of −1 to −3 μm. Micrographs were collected with a pixel size of 1.07 Å. The total dose, 60 e-·Å^-2^, was acquired by using a dose rate of 8.54 e-·pixel^-1^·s^-1^ (7.49 e-·Å^-2^·s^-1^) across 40 frames for 8 s total exposure time.

### Image processing

An overview of the image processing protocol is shown in **Supplementary Figure 11**. All processing was completed in Relion-3.0.7 unless stated otherwise.^44^ The initial drift and beam induced motion was corrected for using MotionCor2, and Ctf estimation was performed using Gctf. Automated particle picking was performed in crYOLO using the general model. For the Pico145-bound structure, this resulted in a particle stack containing 612,983 particles which was imported into Relion-3.0.7. An initial round of 2D classification resulted in a particle stack consisting of 519,962 particles. A 3D initial model was generated in Relion and low-pass filtered to 60 Å when used in 3D classification. The best class, consisting of 266,887 particles, was taken forward and refined with C4 symmetry imposed to give a global resolution of 3.3 Å after post-processing, with resolutions estimated by the gold standard FSC = 0.143 criterion. Three iterative rounds of CTF refinement and particle polishing were completed, which improved the resolution of the map to 3.1 Å. The polished particles underwent two further rounds of 3D classification resulting in a particle stack containing 158,111 particles which was refined to 3.0 Å. A local resolution map was calculated in Relion which showed that the core of the structure was at a higher resolution (2.8 Å) than the global average. For the structure in the presence of ZnCl_2_, crYOLO picked 552,075 particles, which was reduced to 479,810 upon 2D classification. 3D classification further reduced this to 228,615 particles, which were refined and post-processed, resulting in a 3.1 Å structure. After three iterative rounds of CTF refinement and particle polishing, the resolution of the map was improved to 2.8 Å – further 3D classification did not improve the map. For the 50 μM (partial occupancy) Pico145 structure, micrographs were subjected to crYOLO particle picking (536,348 particles), followed by 2D classification (509,928 particles), 3D classification (158,548 particles). 3D classified particles were subjected to refinement and post-processing, resulting in a 3.1 Å structure. Three rounds of CTF refinement and particle polishing improved the overall resolution to 2.9 Å – further 3D classification did not improve the map.

### Creation of PDB file

The model of TRPC5 with 100 μM Pico145 was produced by manual fitting of 6aei into the model. Several rounds of real space refinement in Phenix were performed before fitting Pico145 into the map in COOT.

### Docking studies

The region of the TRPC5 tetrameric structure assigned for docking studies was chosen based upon residues identified to be close Pico145 in our TRPC5:Pico145 structure (Y524, L572, Q573, F576, W577, V579, F599, Y603 and V614). A 25 Å clip of the TRPC5 structure around these residues was termed as the receptor for docking studies using Glide (Schrödinger Release 2019-4, Glide, Schrödinger, LLC, New York, NY, 2020).^45^ The TRPC5 pdb file was prepared using the Protein Preparation Wizard in the Maestro Graphical User Interface (GUI). This aimed to remove any steric clashes of amino acid side chains and optimise the position of hydrogen atoms to facilitate docking studies. The receptor grid was generated using Schrödinger software, allowing docking of ligands in a 36×36×36 Å grid. The Pico145 ligand was prepared using the OMEGA module^46^ of OpenEye software (OMEGA version 2.5.1.4 OpenEye Scientific Software, Santa Fe, NM; http://www.eyesopen.com) to produce an energy-minimised 3D structure before importing into the Maestro GUI. Docking of Pico145 was carried out using the Glide module of Schrödinger software using the XP mode with flexible ligand sampling and biased sampling of torsions for all predefined functional groups. Epik state penalties were added to the docking score. A maximum of 10 poses for the ligand were requested in the output file and post-docking minimisation was carried out.

### Intracellular Ca^2+^ measurements

Intracellular calcium ([Ca^2+^]_i_) was measured using the ratiometric Ca^2+^ dye Fura-2. Cells were plated in 6-well plates at 1 million cells per well and the following day were transfected with 2 μg DNA using 6 μl of JetPrime (VWR International) according to the manufacturer’s protocol. The next day cells were washed, trypsinised and plated onto black, clear bottom poly-D-lysine coated 96-well plates with 60,000 cells per well. 24 h after plating, the cells were loaded with the Fura-2 dye by removal of media and addition of SBS (NaCl 130 mM, KCl 5 mM, glucose 8 mM, HEPES 10 mM, MgCl_2_ 1.2 mM, and CaCl_2_ 1.5 mM; all supplied by Sigma Aldrich) containing 2 μM Fura-2 acetoxymethyl ester (Fura-2 AM; ThermoFisher Scientific) and 0.01% pluronic acid (Merck) for 1 h at 37 °C. After this incubation, cells were washed with fresh SBS and incubated at room temperature for a further 30 min. SBS was then changed to recording buffer (SBS with 0.01% pluronic acid and 0.1% DMSO to match compound buffer) immediately prior to experimentation. In the case of inhibitor studies the buffer was replaced with SBS with 0.01% pluronic acid and the relevant inhibitor or vehicle. Cells were then incubated for 30 min prior to the experiment. Measurements were carried out using a FlexStation (Molecular Devices, San Jose, CA), using excitation wavelengths of 340 and 380 nm, at an emission wavelength of 510 nm. ([Ca^2+^]_i_ recordings were performed at room temperature at 5 s intervals for 300 s (unless stated otherwise). EA or AM237 was added by the FlexStation from a compound plate (final DMSO concentration 0.1%) after recording for 60 s. Responses were calculated at 240-300 s compared to the signal at baseline (0-60 s), and an average was taken within each individual experiment. Averages from individual experiments were normalised relative to the maximum signal for each TRPC5 variant/activator combination, combined, and fit (using a variable slope, 4-parameter fit) with GraphPad Prism 8 with [agonist] for EA or AM237, or [inhibitor] for Pico145 vs response.

### Chemicals

Pico145^23^ and AM237^30^ were prepared according to previously reported procedures. (-)-englerin A (EA) was obtained from PhytoLab (Vestenbergsgreuth, Germany). AM237, Pico145 and EA were made up as 10 mM stocks in 100% DMSO, aliquots of which were stored at −20 °C (AM237 and Pico145) or −80 °C (EA). Further dilutions of compounds were made in DMSO and these were dissolved 1:1000 in compound buffer (SBS + 0.01% pluronic acid) before being added to cells. Fura-2 AM (Invitrogen UK) was dissolved at 1 mM in DMSO.

## Acknowledgements

This work was supported by the BBSRC (BB/P020208/1). Professor Eric Gouaux (Vollum Institute) is kindly acknowledged for the BacMam vector. Large-scale tissue culture was performed in the University of Leeds Protein Production Facility (funded by the University of Leeds and the Royal Society), with support provided by Dr Brian Jackson. We thank the Astbury Biostructure Laboratory (funded by the University of Leeds and the Wellcome Trust) for support of electron microscopy work, Negative stain microscopy was performed using a T12 microscope funded by the Wellcome Trust (090932/Z/09/Z). Cryo-EM was performed with a Titan Krios microscope with an energy filtered Gatan K2 XP summit direct electron detector (108466/Z/15/Z).

## Author contributions

DJW performed the design and cloning of constructs, intracellular calcium measurements and protein purification. DJW, RMJ, and SPM performed electron microscopy studies, including data analysis and model building. KJS performed docking studies. DJW, KJS, RMJ, SPM and RSB analysed data. DJB, SPM and RSB conceived the project and generated research funds. SPM and RSB led the project. DJW, KJS, SPM and RSB prepared figures and wrote the manuscript. All authors commented on the manuscript.

## Additional Information

**Supplementary Figures 1-11** and **Supplementary Tables 1-2** are attached below.

### Supplementary Figures

**Supplementary Figure 1:**
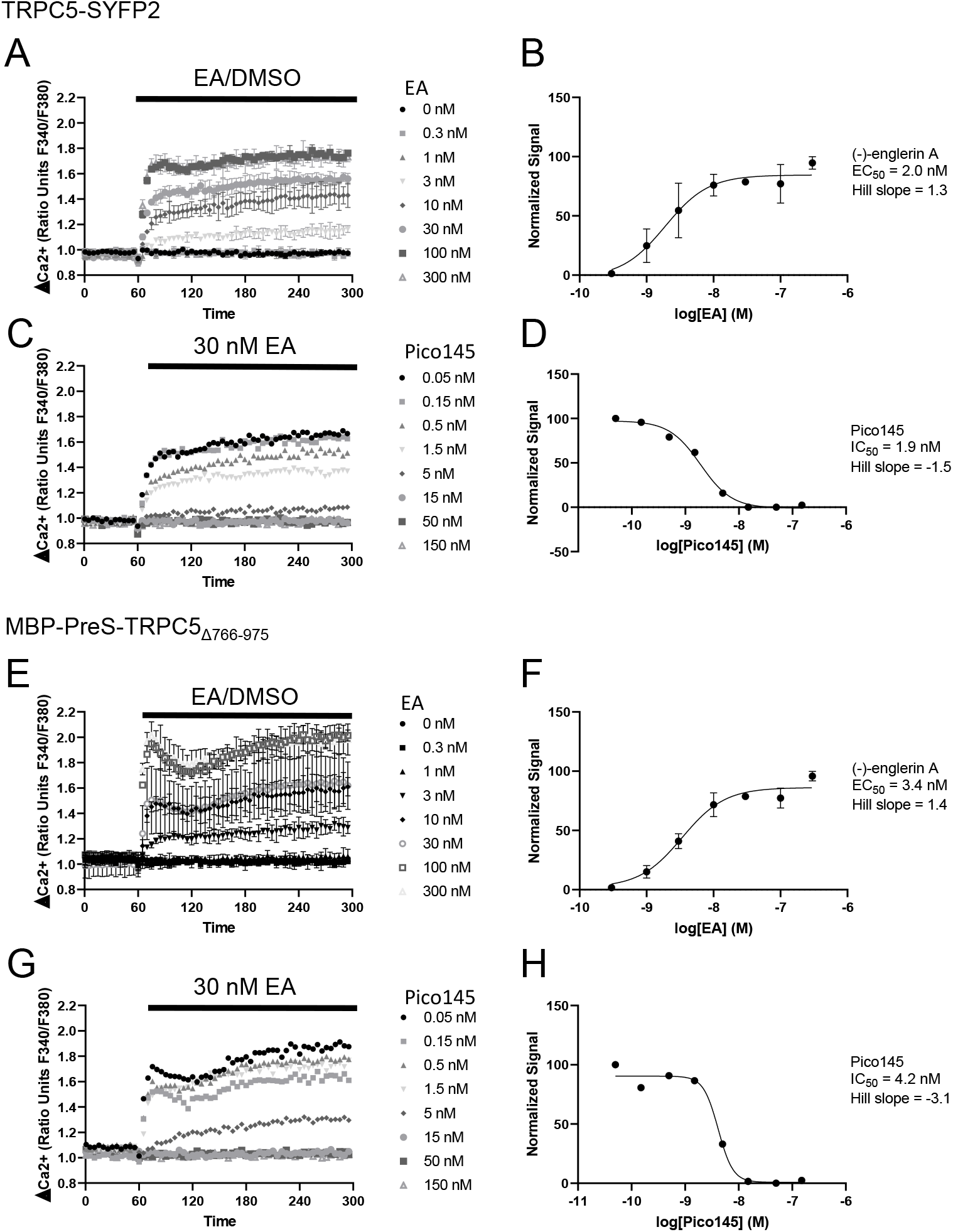
Response of C-terminally truncated, N-terminally labelled hTRPC5 to EA and Pico145 is comparable with that of full-length hTRPC5-SYFP2. A,E) Representative traces from single 96-well plates (N = 3) showing an increase in [Ca^2+^]_i_ in response to 0.32 nM to 5 μM EA of HEK 293 cells transiently expressing TRPC5-SYFP (A) or MBP-PreS-TRPC5_Δ766-975_ (E). B,F) Concentration-response data for experiments (A) and (E) (mean normalised response ± SEM; n/N = 3/8). C,G) Traces from single 96-well plates (N = 1) showing that the increase in [Ca^2+^]_i_ in response to 30 nM EA is concentration-dependently inhibited by pre-incubation with 0.05 nM to 150 nM Pico145 in HEK 293 cells transiently expressing TRPC5-SYFP (C) or MBP-PreS-TRPC5_Δ766-975_ (G). F,H) Concentration-response data for experiments in (C) and (G). Responses were calculated at 240-300 s compared to [Ca^2+^]_i_ at baseline (0–60 s). Averages of technical repeats (where relevant) of each individual experiment were normalised to the maximum response for each set of experiments, combined, and fit with GraphPad Prism 8 (variable slope, four-parameters fit).

**Supplementary Figure 2:**
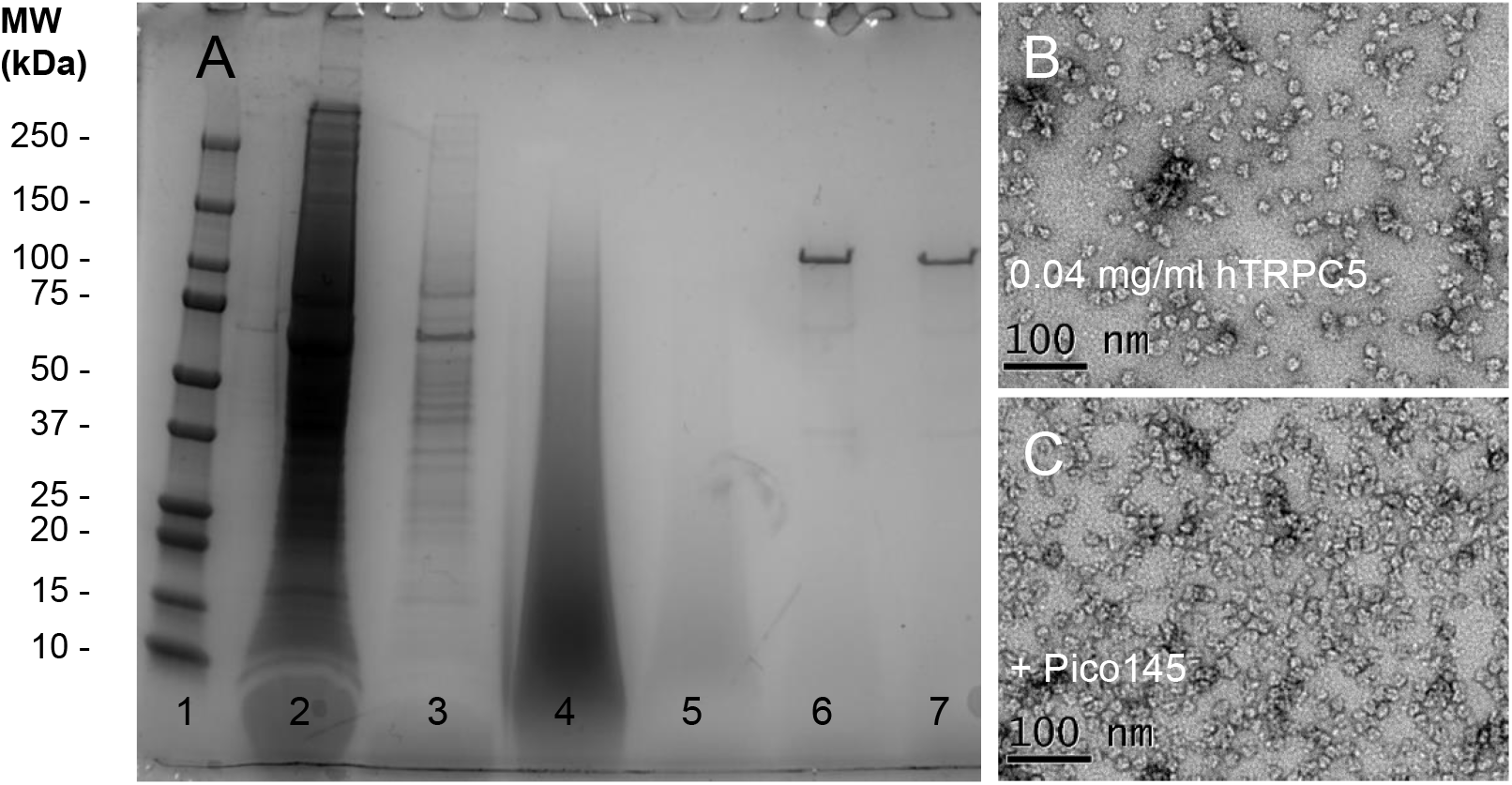
Purification of MBP-PreS-TRPC5_Δ766-975_ and negative stain electron microscopy. A) Purification of hTRPC5 using amylose resin and exchange into PMAL-C8 amphipol. Lanes: **1.** Ladder; **2.** Flow-through A; **3.** Wash A; **4.** Flow-through after Amphipol exchange; **5.** Wash B; **6.** Elution; **7.** Supernatant of ultracentrifugation. B) Negative stain electron microscopy of purified hTRPC5 at 49,000 × magnification. C) Negative stain electron microscopy of purified hTRPC5 (0.04 mg·ml^-1^) at 49,000 × magnification, plus 4 μM Pico145.

**Supplementary Figure 3:**
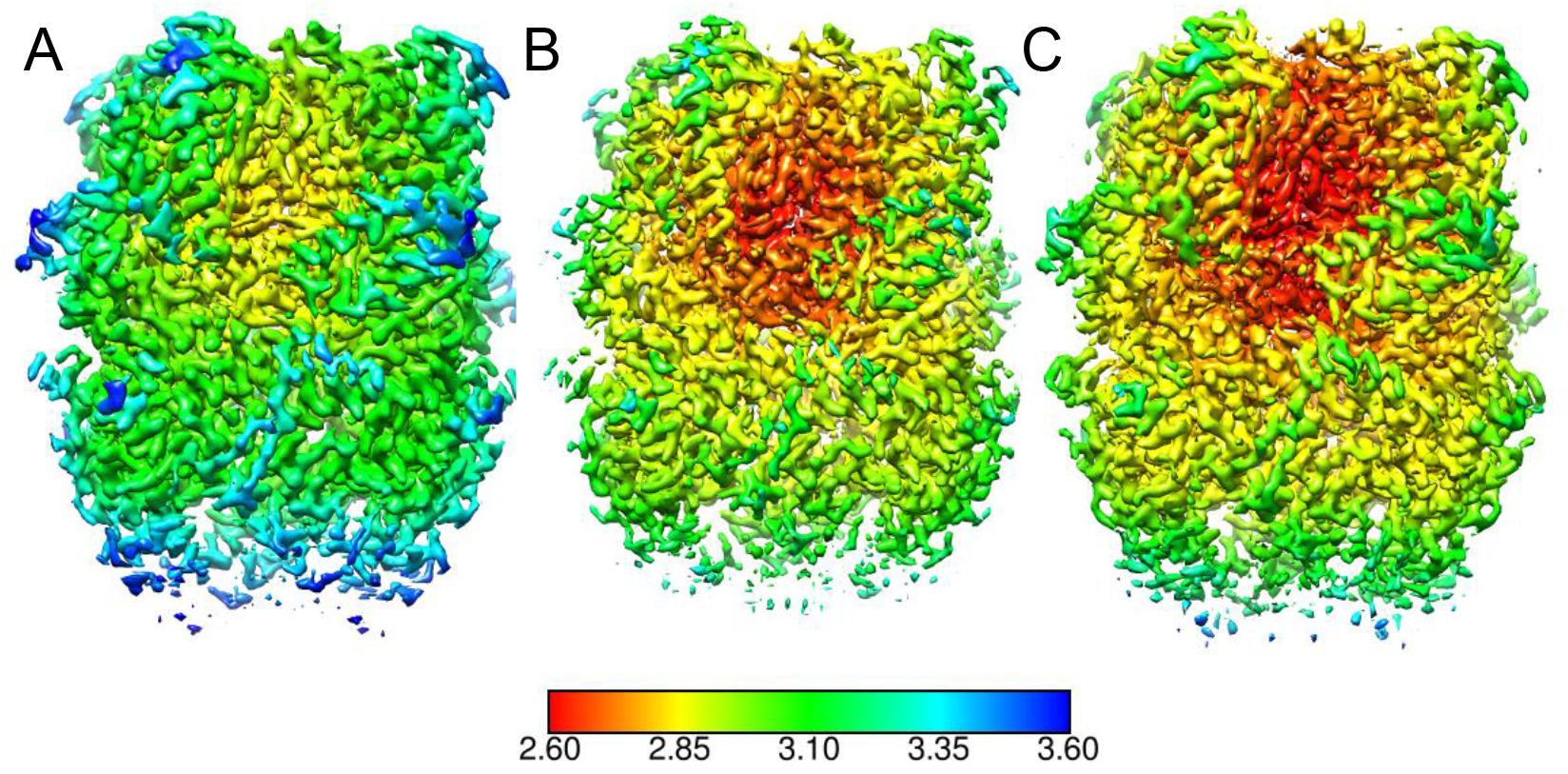
Local resolution of TRPC5 structures. A) Local resolution of the hTRPC5:Pico145 structure. B) Local resolution of the partially Pico145-occupied hTRPC5 structure. C) Local resolution of the hTRPC5 structure in the presence of 20 μM ZnCl_2_.

**Supplementary Figure 4:**
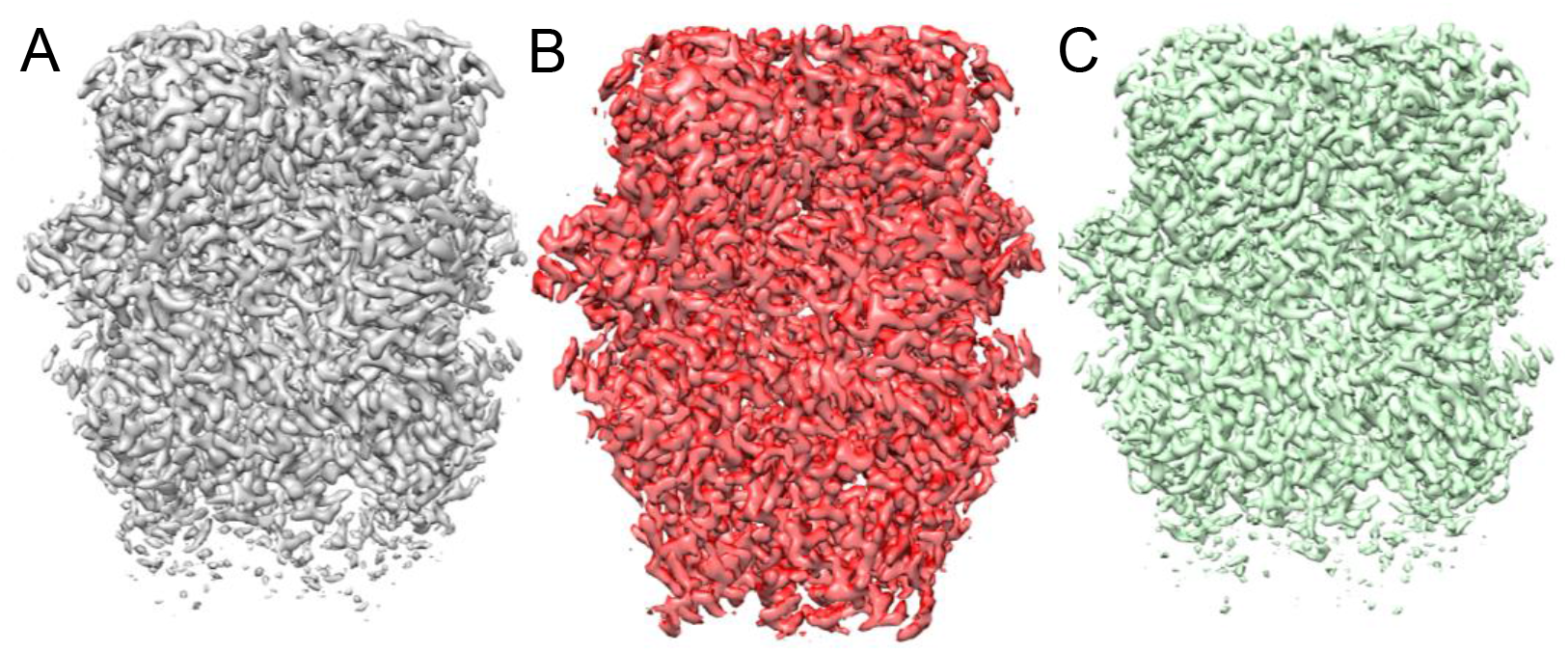
Comparison of overall TRPC5 structures. A) Electron density of hTRPC5 in the presence of 100 μM Pico145. B) Electron density of mTRPC5 *apo* (PDB 6aei; EMDB 9615). C) Electron density of hTRPC5 structure in the presence of 20 μM ZnCl_2_.

**Supplementary Figure 5:**
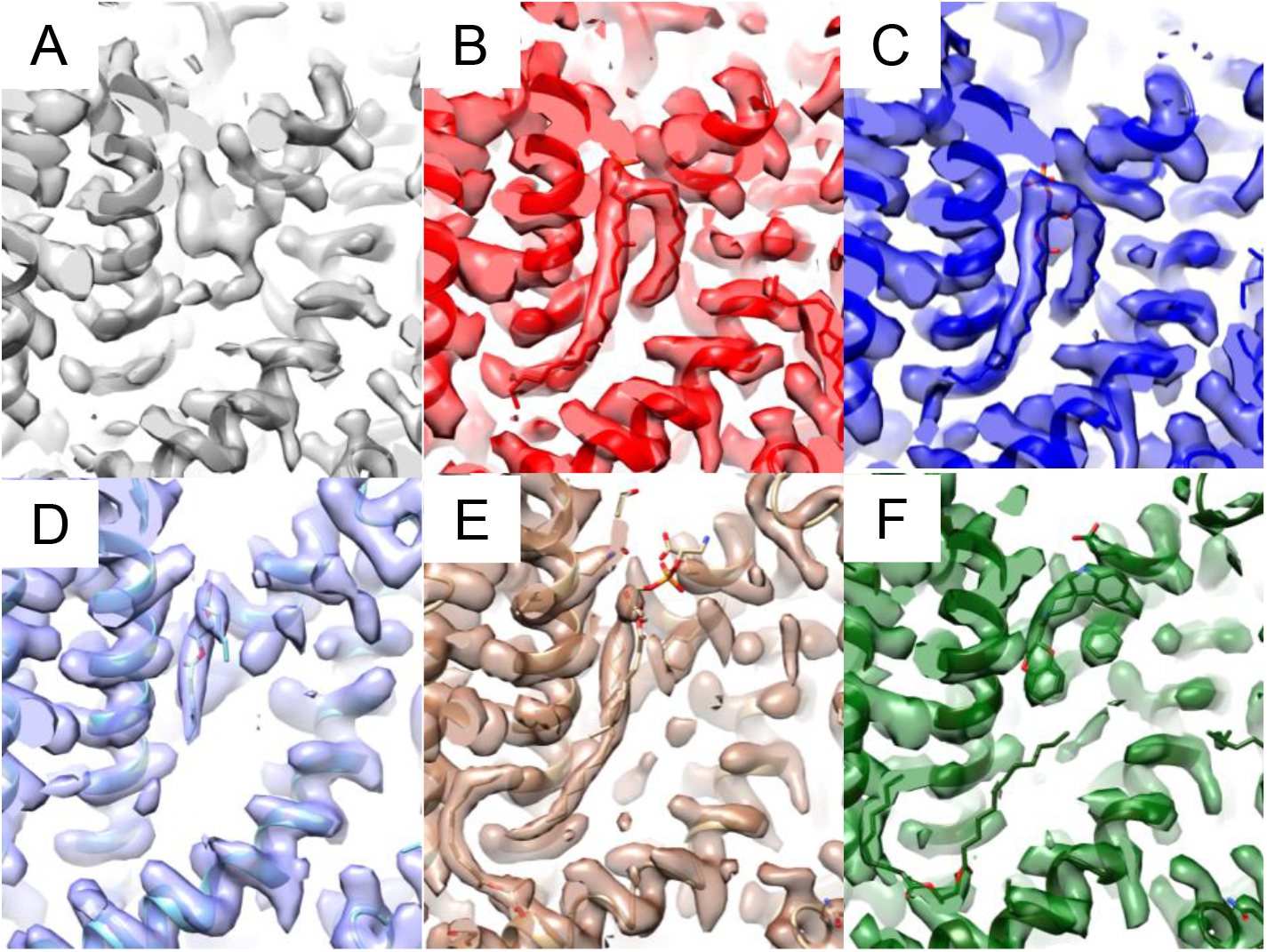
Pico145 binds to a conserved TRPC4/5 lipid binding site. A) Density observed in Pico145-bound hTRPC5 structure (100 μM Pico145). B) Equivalent density in the published mTRPC5 *apo* structure (PDB 6aei; EMDB 9615) that was modelled as a phospholipid. C) The same site from the published mTRPC4 structure (PDB 5z96; EMDB 6901). D) Lipid bound to equivalent residues in the hTRPC3 structure (PDB 6cud; EMDB 7620). E) The equivalent site from the structure of hTRPC6 in complex with the TRPC6 antagonist AM-1473 (PDB 6uz8; EMDB 20953) revealing a bound lipid. F) The same site from the published structure of hTRPC6 in complex with the TRPC6 agonist AM-0883 (PDB 6uza; EMDB 20954), showing the displacement of lipid by the small molecule AM-0883.

**Supplementary Figure 6:**
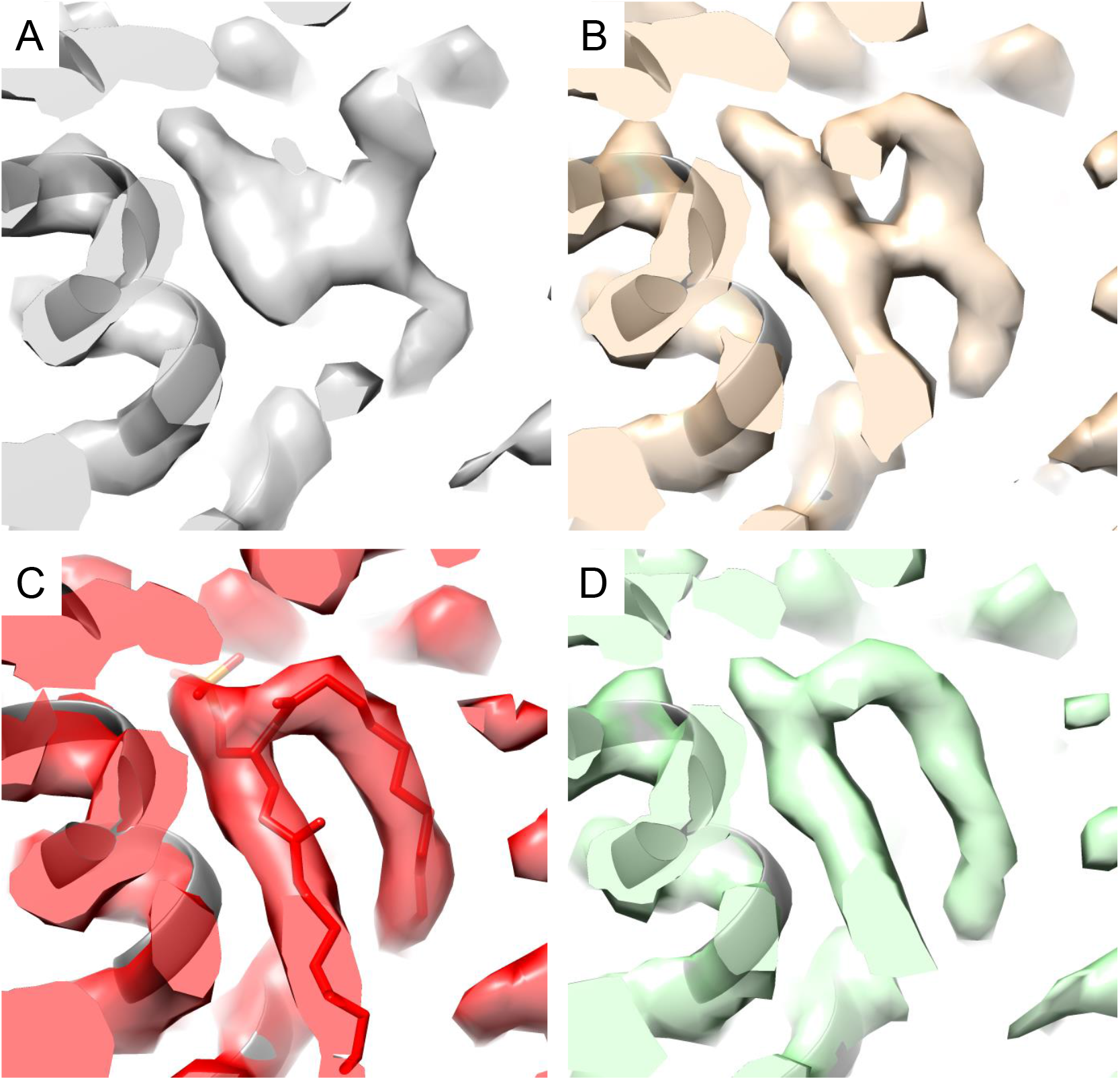
Comparison of the Pico145/lipid binding site of TRPC5 in three structures. A) Lipid/xanthine binding site of hTRPC5 in the presence of 100 μM Pico145. B) Lipid/xanthine binding site of hTRPC5 in the presence of 50 μM Pico145. C) Lipid/xanthine binding site of mTRPC5 *apo* (PDB 6aei; EMDB 9615). D) Lipid/xanthine binding site of hTRPC5 in the presence of20 μM ZnCl_2_.

**Supplementary Figure 7:**
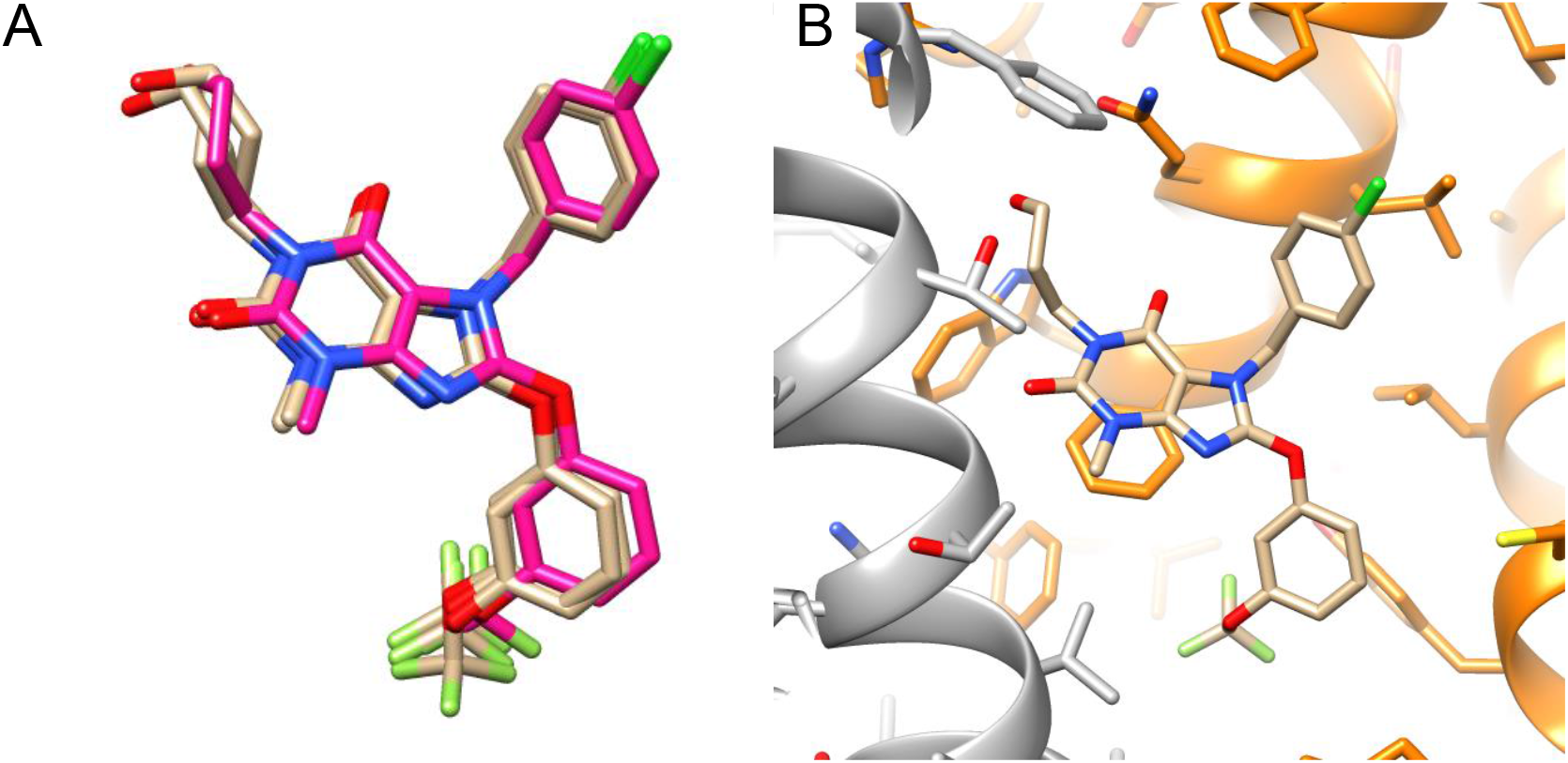
Docking of Pico145 into hTRPC5 shows key interactions. A) The top three docked poses of Pico145 in TRPC5 (peach) overlaid with the refined structure (magenta) in the absence of the protein structure. There is good agreement between the structures and the position of the xanthine core is highly conserved. The different positions of the flexible 3-hydroxypropyl substituent at N-1 suggest it is flexible within the TRPC5 binding pocket. B) The best scoring docking pose is shown relative to the PDB file used for docking (as in Figure 3).

**Supplementary Figure 8:**
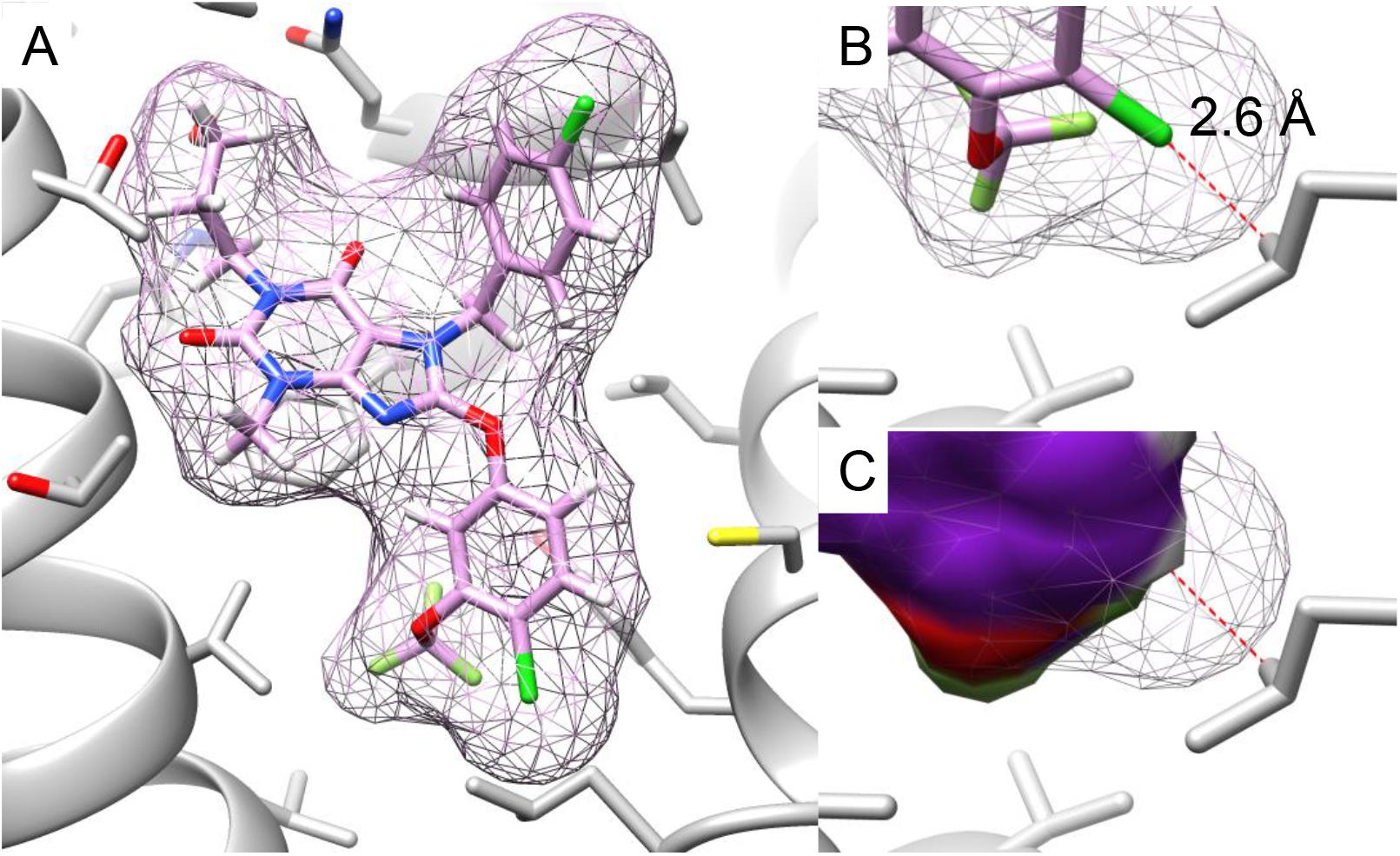
AM237 in the same pose as Pico145 in TRPC5 would clash with L521. AM237 was modelled into the top scoring docking pose of Pico145 in TRPC5. A) The full AM237 structure is shown in mesh format. B) A zoomed in view of the clash between chlorine atom and L521. C) The same view as in (B), but with the surface of docked Pico145 structure shown as solid colour, showing the removal of the clash with L521.

**Supplementary Figure 9:**
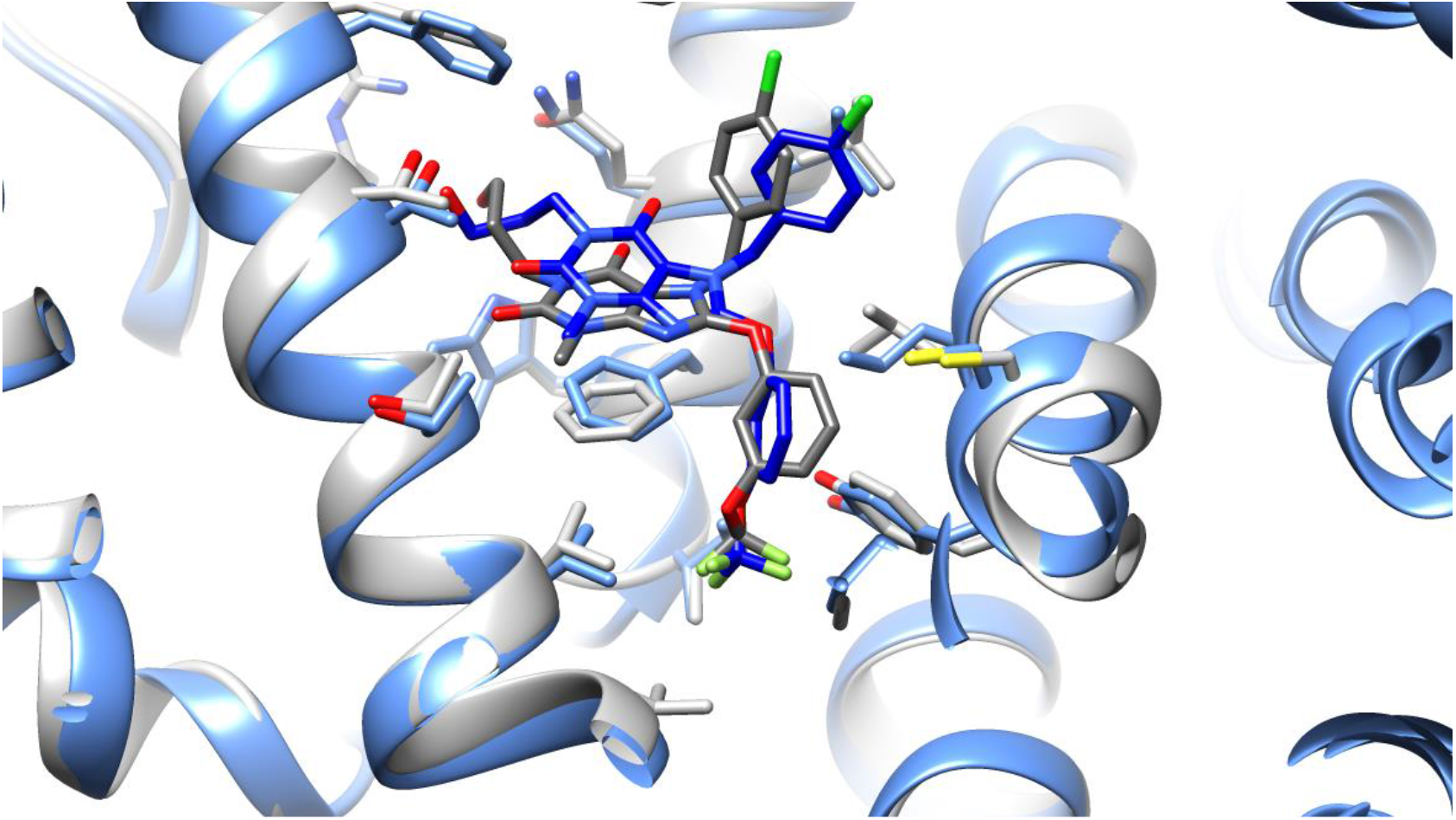
The docked poses of Pico145 in TRPC5 and TRPC4 are similar. Pico145 was docked into TRPC5 (grey: our TRPC5:Pcio145 structure minus refined Pico145) and TRPC4 (PDB 5z96; blue). All interacting residues are conserved (**Supplementary Table 2**) except V579 and the docking poses overlay well. Only the top scoring docking pose (of three) is shown for each TRPC structure.

**Supplementary Figure 10:**
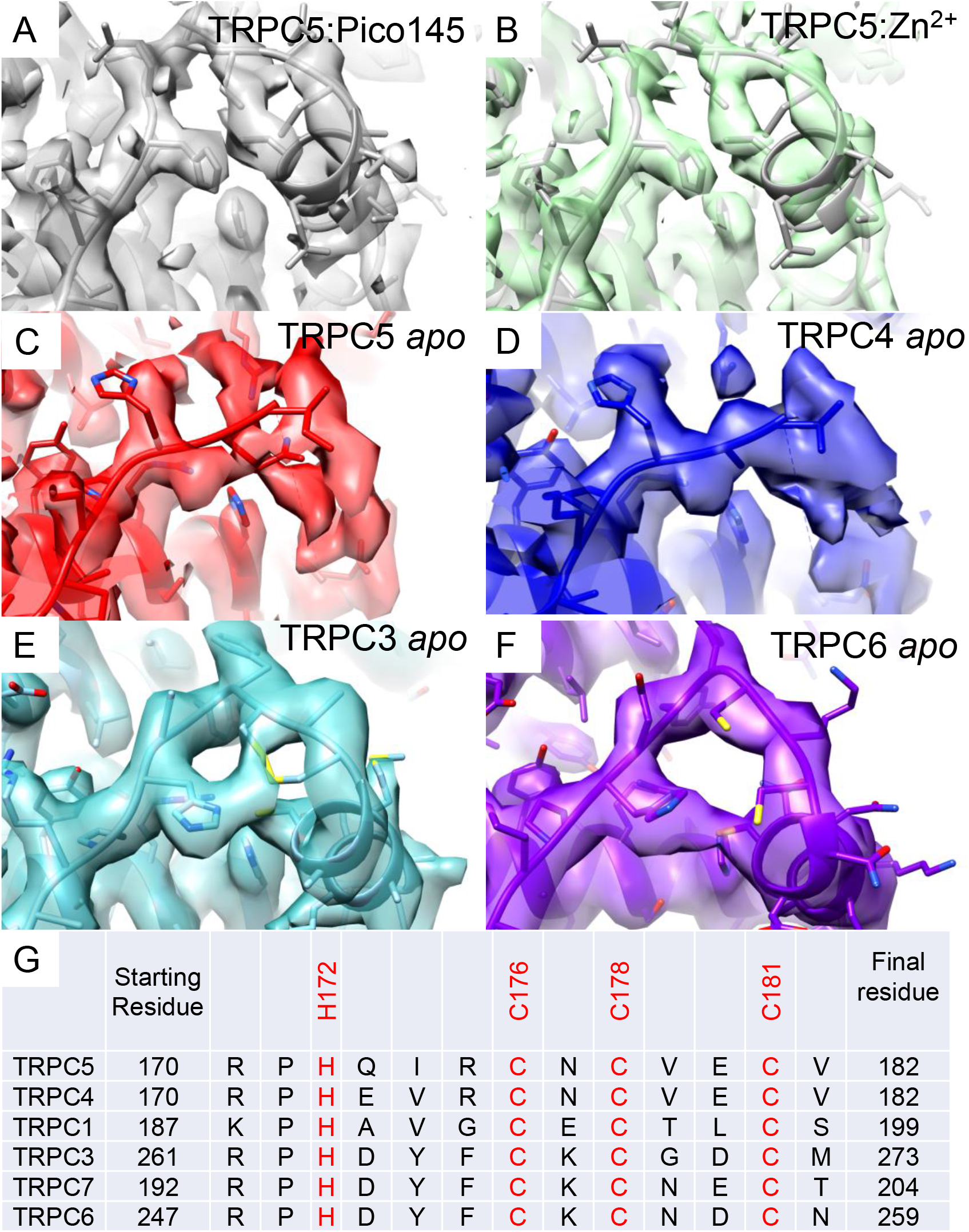
Alignment of the putative zinc binding sites of TRPC channels. A-F) The putative zinc binding site of hTRPC5 in the presence of 2 μM Pico145 (A) and hTRPC5 in the presence of 20 μM ZnCl_2_ (B), mTRPC5 (PDB 6aei; EMDB 9615; (C)), mTRPC4 (PDB 5z96; EMDB 6901; (D)), hTRPC3 (PDB 6cud; EMDB 7620; (E)) and hTRPC6 (PDB 5yx9; EMDB 6856; (F)) is shown with PDB files fit into experimental densities. G) Sequence alignment of hTRPC5 with the other human TRPC proteins, showing conservation of His and Cys residues in the putative zinc binding sites.

**Supplementary Figure 11:**
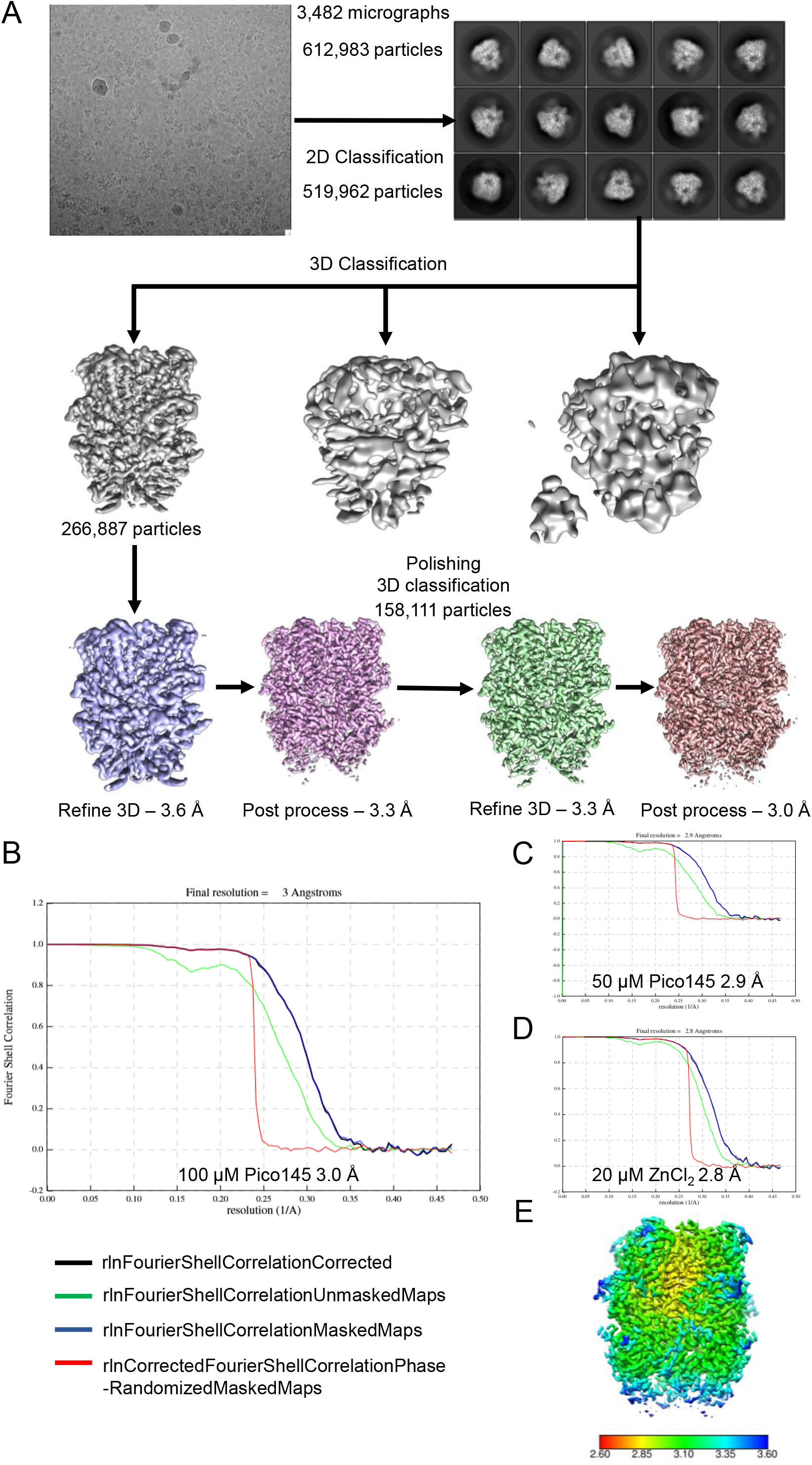
TRPC5 Cryo-EM data processing. A) Representative micrograph of grids prepared from 2.0 mg·ml^-1^ MBP-PreS-hTRPC5_Δ766-975_ with 100 μM Pico145, and graphical representation of 2D and 3D classification, refinement and post-processing as described in the Methods section. B) Fourier shell correlation (FSC) curves for the 3D reconstruction from grids prepared from 2.0 mg·ml^-1^ of MBP-PreS-hTRPC5_Δ766-975_ with 100 μM Pico145 (3.0 Å resolution). C) FSC curves for the 3D reconstruction from grids prepared from 1.0 mg·ml^-1^ MBP-PreS-hTRPC5_Δ766-975_ with 50 μM Pico145 (2.9 Å resolution). D) FSC curves for the 3D reconstruction from grids prepared from 1.0 mg·ml^-1^ MBP-PreS-hTRPC5_Δ766-975_ with 20 μM ZnCl2 (2.8 Å resolution). E) Local resolution estimation from Relion.

### Supplementary Tables

**Supplementary Table 1:**
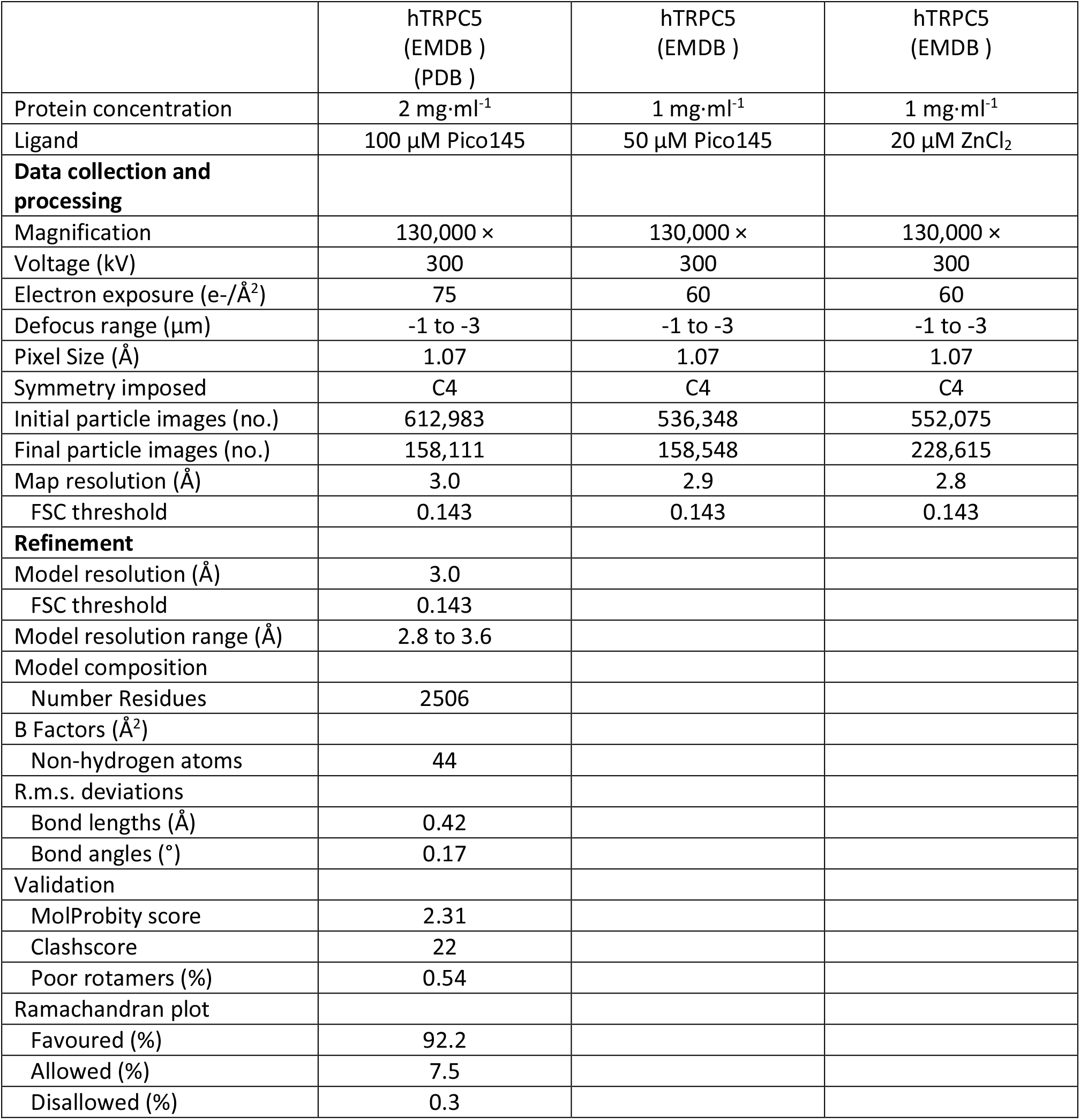
Cryo-EM data collection, refinement and validation statistics.

|  |  |  |  |
| --- | --- | --- | --- |
|  | hTRPC5<br>(EMDB )<br>(PDB ) | hTRPC5<br>(EMDB ) | hTRPC5<br>(EMDB ) |
| Protein concentration | 2 mg·ml<sup>-1</sup> | 1 mg·ml<sup>-1</sup> | 1 mg·ml<sup>-1</sup> |
| Ligand | 100 µM Pico145 | 50 µM Pico145 | 20 µM ZnCl<sub>2</sub> |
| **Data collection and processing** |  |  |  |
| Magnification | 130,000 × | 130,000 × | 130,000 × |
| Voltage (kV) | 300 | 300 | 300 |
| Electron exposure (e-/Å<sup>2</sup>) | 75 | 60 | 60 |
| Defocus range (µm) | -1 to -3 | -1 to -3 | -1 to -3 |
| Pixel Size (Å) | 1.07 | 1.07 | 1.07 |
| Symmetry imposed | C4 | C4 | C4 |
| Initial particle images (no.) | 612,983 | 536,348 | 552,075 |
| Final particle images (no.) | 158,111 | 158,548 | 228,615 |
| Map resolution (Å) | 3.0 | 2.9 | 2.8 |
| FSC threshold | 0.143 | 0.143 | 0.143 |
| **Refinement** |  |  |  |
| Model resolution (Å) | 3.0 |  |  |
| FSC threshold | 0.143 |  |  |
| Model resolution range (Å) | 2.8 to 3.6 |  |  |
| Model composition |  |  |  |
| Number Residues | 2506 |  |  |
| B Factors (Å<sup>2</sup>) |  |  |  |
| Non-hydrogen atoms | 44 |  |  |
| R.m.s. deviations |  |  |  |
| Bond lengths (Å) | 0.42 |  |  |
| Bond angles (°) | 0.17 |  |  |
| Validation |  |  |  |
| MolProbity score | 2.31 |  |  |
| Clashscore | 22 |  |  |
| Poor rotamers (%) | 0.54 |  |  |
| Ramachandran plot |  |  |  |
| Favoured (%) | 92.2 |  |  |
| Allowed (%) | 7.5 |  |  |
| Disallowed (%) | 0.3 |  |  |

**Supplementary Table 2:**
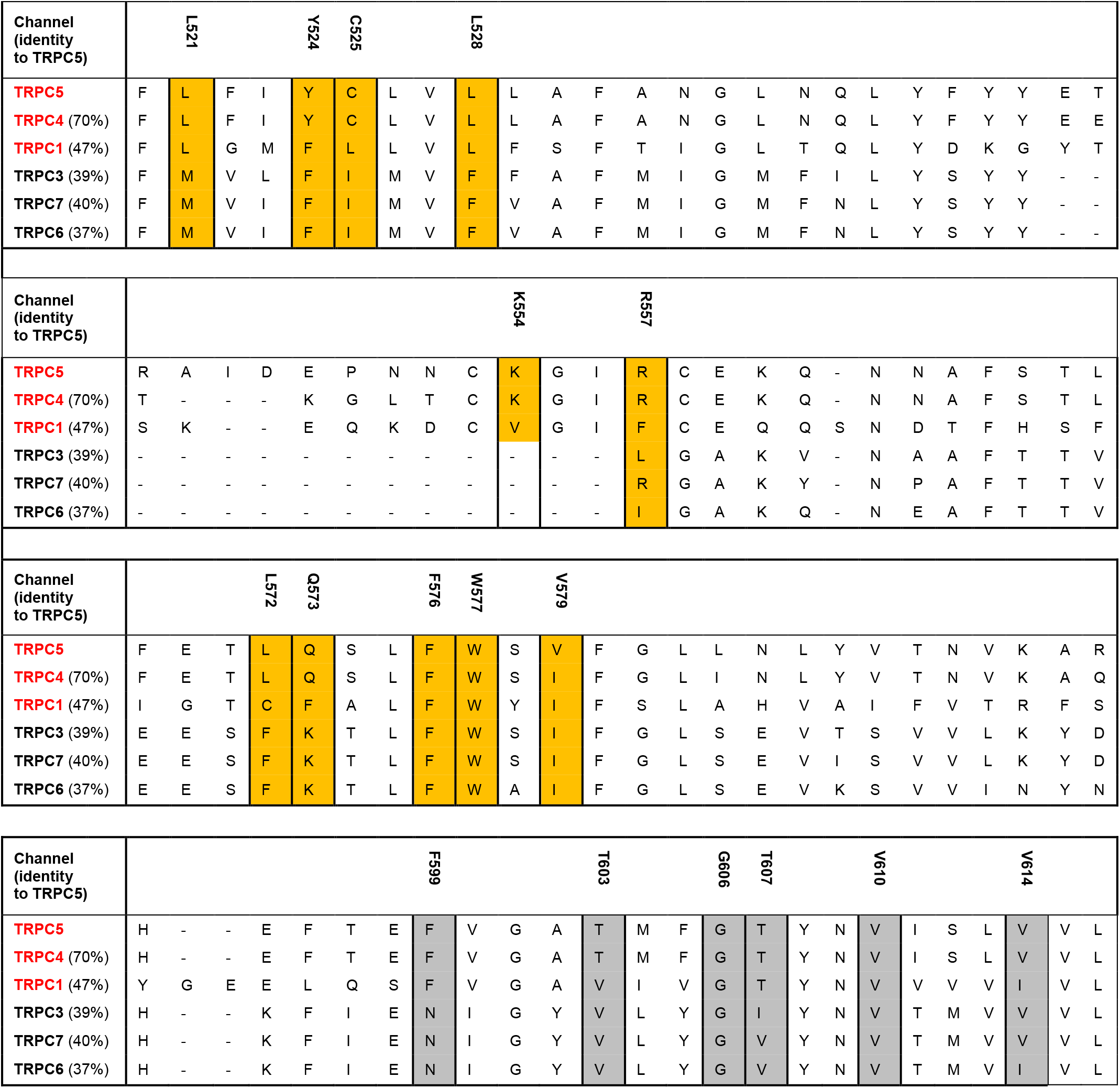
Sequence alignment of xanthine binding site within the TRPC family. Residues 520-616 of TRPC5 were aligned within the human TRPC proteins. Overall identity to TRPC5 is shown and residues equivalent to those close to Pico145 in the TRPC5 structure are shaded in orange (monomer 1; **Figure 3A**) and grey (monomer 2). The numbers above residues above are those in TRPC5. TRPC1, TRPC4 and TRPC5 are coloured in red because channels containing these monomers can be modulated by xanthines such as Pico145 and AM237.

